# Deep-Pose-Tracker: an automated behavioural analysis framework for *Caenorhabditis elegans*

**DOI:** 10.1101/2025.11.23.689997

**Authors:** Debasish Saha, Shivam Chaudhary, Dhyey Vyas, Anindya Ghosh-Roy, Rati Sharma

## Abstract

Tracking and analyzing animal behaviour is a crucial step in fields such as neuroscience and developmental biology. Behavioural studies in the nematode *C. elegans*, for example, help in understanding how organisms respond to external cues and how the specific physiological responses link to either instantaneous or learned behaviours. Although tracking behaviour through locomotion patterns and postural dynamics is routine, it becomes laborious and time-consuming when performed manually. Automation of this process is therefore crucial for accurate and fast detection and analysis. To this end, we report Deep-Pose-Tracker (DPT), a YOLO (You Only Look Once)-based model for automated pose detection of *C. elegans* from videos and images. The module is further utilized for several downstream analysis algorithms to quantify essential behavioural features, including locomotion speed, orientation, forward or reverse locomotion, and complex body bends such as omega turns. In addition, it includes eigenworms decomposition to represent complex posture dynamics in a low-dimensional space. The model shows reliable performance on the validation and test datasets, with high inference speed, while being user-friendly. DPT, therefore, can be a valuable toolkit for automated behavioural quantification of *C. elegans* under varying experimental stimuli.

## I. INTRODUCTION

Behavioral studies are of fundamental importance across a wide range of research fields, including neuroscience, genetics, ecology, evolutionary biology, and drug discovery. They provide the first preliminary insight into how a stimulant can translate into a quantifiable physiological response, such as a change in a movement pattern [1, 2]. The measurable changes in behaviour can then lead to a better understanding of the kind of strategies animals employ for both survival and adaptation in changing environments [3–8]. The nematode (roundworm), *Caenorhabditis elegans*, is one such well-established model organism where behaviour is studied extensively and correlates well with stress response, ageing, neuronal signalling, and survival under different external conditions. Its short lifespan (3-4 weeks), short generation time (~2 days) and fully mapped nervous system consisting of 302 interconnected neurons make it suitable for controlled laboratory studies, where specific neuronal activity can be interpreted through physiological responses [9]. *C. elegans*, therefore, presents a good system for behavioural studies that is primed to benefit greatly from automation.

Behavioural studies in most animals, including *C. elegans*, typically involve observing specific animal responses such as locomotion patterns, speed, body shape variations, and orientation under various experimental conditions. Quantification of these parameters requires automated tools that enable robust comparisons of typical, learned, and atypical behaviours across conditions. One of the most common features that benefit from automation is tracking of animal trajectories [10–12]. In the context of *C. elegans*, tracking methods in the past relied on traditional image processing techniques, in which worms are first separated from the background using a thresholding technique, creating a mask around the worm. The mask is then used for tracking and skeletonization to estimate the worm body posture [13–16]. These classical approaches, though, require manual parameter tuning, where performance may be found to degrade in noisy, low-contrast or uneven imaging conditions. Hardware-based trackers like Track-a-Worm and Track-a-Worm2.0 were designed, on the other hand, to analyse worm movement and posture on automated movable microscopy stages where worms are kept centrally located under the camera [16–22]. For high-throughput experiments, Multi-Worm Tracker (MWT) was developed, which could track up to 120 worms simultaneously [23]. More recently, Tierpsy Tracker has emerged as a popular tool due to its effectiveness as well as its ability to extract interpretable locomotion features [24, 25]. In addition to these, there exist several open-source programs like ImageJ (with TrackMate), which enable threshold-based multiple worm tracking [26]. The classical tracking methods work well in controlled and uniform imaging conditions with appropriate parameter tuning; however, their performance may degrade in challenging scenarios, such as crowded environments, overlapping individuals, self-occlusions and noisy backgrounds.

Deep learning (DL) models, on the other hand, employ a fundamentally different approach, in which the neural networks are trained to learn complex patterns directly from the input data and generate accurate predictions. Unlike classical threshold-based techniques, DL models are generally more robust to noisy, low-contrast, and uneven imaging conditions, thereby reducing the requirement for manual parameter tuning and enabling accurate tracking and pose estimation from images and videos [27–37]. Recently, a U-Net-based pose estimation model demonstrated strong performance on real, low-resolution, and noisy images, despite being trained on synthetic worm-posture images [35]. Other DL-based pose estimation frameworks, such as Wormpose [30], DeepTangle [32], and DeepTangleCrawl [36], are built upon CNN (convolutional neural network) architectures and are trained using synthetic images or a combination of synthetic and manually annotated images to predict complex body postures like omega and delta turns, even when worms are self-occluded. For segmentation tasks as well, DL models provide accurate pixel-wise segmentation of *C. elegans* including partially occluded body shapes [34, 38]. Several other models, such as LEAP, SLEAP and DeepLabCut, are highly suitable for working with different model organisms and generate high-precision, interpretable behavioural data, again based on DL frameworks [27, 29, 31]. Therefore, DL models offer several advantages for behavioural analyses, particularly due to their robustness to noise, ability to learn and interpret complex patterns, and improved performance in challenging conditions [30, 32, 33, 36].

There exist several models for tracking locomotion and segmentation of *C. elegans*; however, only a few are capable of accurately predicting worm postures (body shape) [24, 30, 32, 33, 35]. Tierpsy tracker and Wormpose are among the widely used frameworks that not only estimate worm postures, but can also identify heads and tails of the worms, which is of significant importance both for neuronal and behavioural studies [24, 30]. Tierpsy tracker relies upon traditional image processing techniques, uses thresholding to subtract the background and generate a mask around the worm, and then predicts body centerline [24]. However, its performance degrades when applied to challenging situations, like overlaps and self-occlusions. In addition, Tierpsy tracker requires manual parameter tuning, which is labour-intensive and sensitive to imaging conditions. Wormpose, on the other hand, is based on a CNN architecture and is highly effective for detecting complex poses, including self-occlusions, without requiring manual parameter tuning [30]. However, Wormpose is primarily designed for single-worm posture detection, thereby, limiting its application for high throughput multi-worm analysis. Furthermore, it performs time-series analysis to identify heads and tails, which requires sufficient body movement. As a result, the head-tail assignment becomes difficult for stationary worms or when applied on isolated images. In addition, unlike Tierpsy tracker, Wormpose does not directly provide any interpretable behavioural features, which limits downstream analyses and quantification of behaviours under varying experimental conditions. Therefore, there remains a need for a more general and automated framework for behavioural quantification. In particular, the model should be capable of detecting complex postures of multiple worms directly from experimental images and videos, while minimizing the need for explicit manual parameter tuning. Additionally, integrating pose estimation with downstream behavioural analyses within a unified framework would facilitate automated quantification of worm locomotion and posture dynamics. Such a system can then become a valuable tool for the *C. elegans* research community.

With this motivation, in this work, we develop Deep-Pose-Tracker (DPT), a deep learning-based framework for multi-worm pose detection and tracking of *C. elegans*. The model is built upon the YOLO (You Only Look Once)^1^ architecture, which was originally introduced in 2016 [39], and has been subsequently improved through multiple upgrades [40–45]. It has become a widely popular computer vision model for object detection, tracking, pose detection, and image segmentation, due to its high inference speed and robust accuracy [39, 43, 46–48]. In particular, YOLO-based models are highly effective for multi-class and multi-object detection tasks, making them suitable for varied and crowded environments. For these reasons, YOLO has been applied extensively across diverse areas, such as traffic monitoring, surveillance, agriculture, industry, robotics, and biological data analysis [49–63], including recent applications to study *C. elegans* behaviours [33, 64–67]. These YOLO models enable a wide range of functionalities, such as object detection and tracking, pose estimation, segmentation and classification within a unified pipeline, making it suitable for quantitative locomotion analysis within a single framework.

The YOLO-based Deep-Pose-Tracker leverages the CNN architecture to learn complex spatial patterns of the objects (worms in this case) from the training images, and enables accurate pose estimation at high inference speeds. The model detects the head and tail of the worms, which is useful for several downstream analyses, as discussed later. *C. elegans* possesses a highly uniform body shape with a few visually distinguishable features, which makes it difficult for many classical models to reliably identify the heads and tails. However, the proposed YOLO-based DPT model can learn distinctive spatial features directly from raw images during training, and identify heads and tails without any explicit post-processing. As a result, the framework can be implemented to both images and videos under varying experimental conditions. In addition to pose estimation and head-tail identification, the DPT model is integrated with several downstream analyses that are essential for behavioural studies of *C. elegans*. These include measurement of speed, eigenworm decomposition (low-dimensional representation of complex posture dynamics) [68], calculation of spatial exploration and quantification of movement orientation during locomotion. The framework is integrated further to detect and quantify complex behavioural patterns, such as forward-reverse motions and omega turns. A major advantage of the proposed framework is its suitability for working with both multi-object and multi-class detection tasks. To demonstrate this, we have additionally performed a simultaneous counting assay of worms and eggs, which is relevant in the study of chemotaxis and egg-laying. The counting framework is tested across a wide range of imaging conditions within the dataset, with varied background colours, heterogeneous populations, and noisy experimental conditions, while eliminating the need for manual tuning.

The rest of the manuscript is organized as follows. In Sec. II, we provide details of the model development, which include experimental details, dataset preparation and splitting, annotation, and training of the model. The results section (Sec. III) is divided into two subsections. In the first part, we report the model performance on the evaluation and test datasets, and in the second part, we demonstrate the integration of DPT for behavioural quantification of *C. elegans*. In Sec IV, we compare our model with other pre-existing models, specifically highlighting the advantages of this YOLO-based method, while also mentioning the areas where DPT can be improved. Finally, we summarize the key results and conclusions in Sec. V.

## II. MATERIALS AND METHODS

In this section, we discuss the individual steps of preparing the model, which include (A) strain preparation and maintenance, (B) video and image acquisition, (C) annotations and dataset preparation, (D) dataset splitting, (E) training and evaluation, and (F) handling collisions and overlaps. A flowchart is shown in Fig. 1, which outlines the primary steps for data collection and model preparation and the subsequent analysis pipeline.

**Figure 1:**
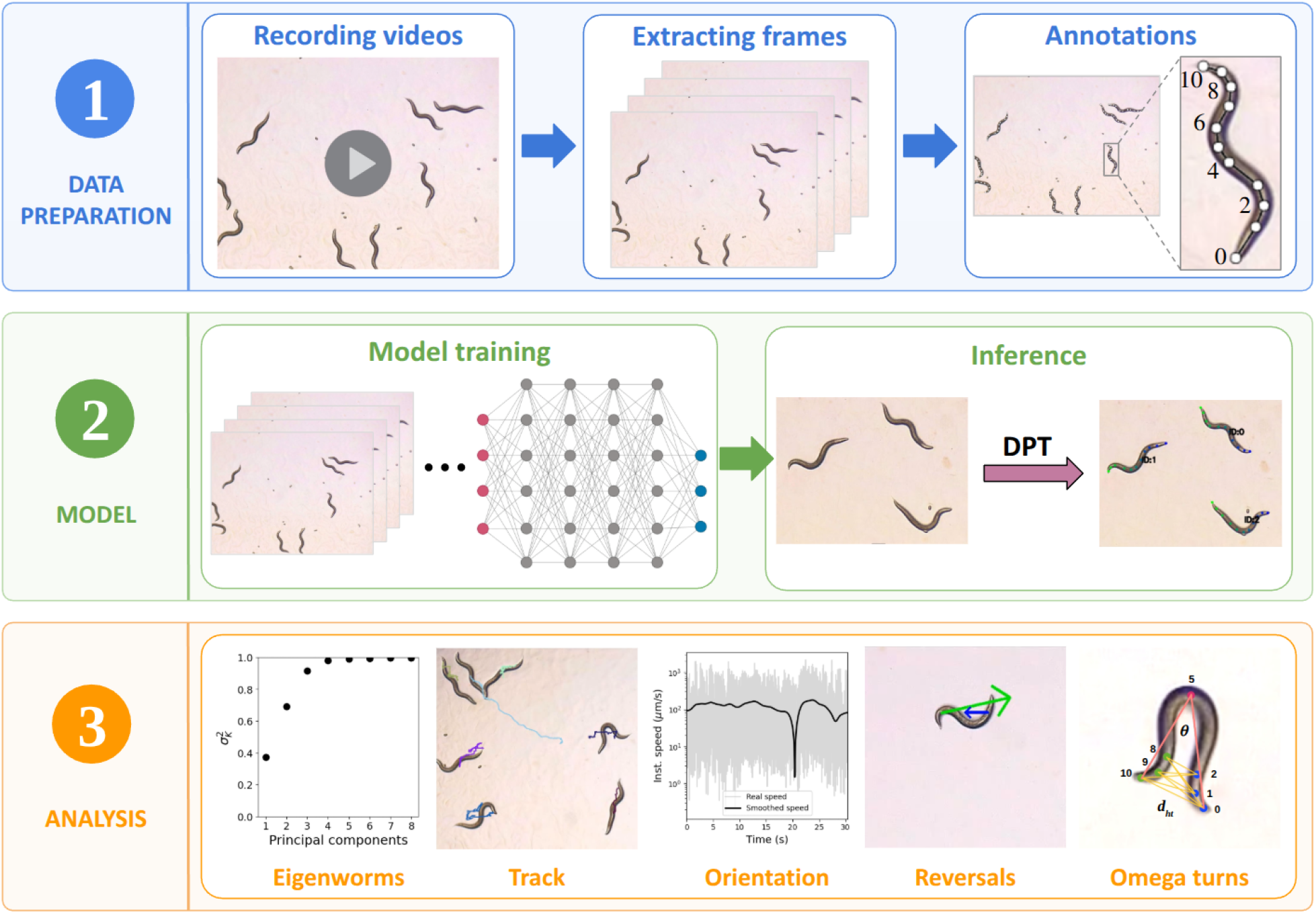
Flowchart of preparing the DPT model. The upper panel shows the dataset preparation step, which is a combination of video capture, frame extraction, and keypoints annotation. In the enlarged box, the keypoints along the body of a worm are shown, starting with 0 assigned to the head and 10 assigned to the tail. The middle panel involves model training and inference. YOLO pose models are trained over the annotated images, and they output keypoint predictions when images are given as input. The third panel highlights the different behavioural quantifications of *C. elegans*, as obtained from the analysis toolkit of DPT.

### A. Strain preparation and maintenance

Day 1 adult *C. elegans* N2 (wild-type) strain was used for imaging, which was further used for model training and behavioural quantification. The worms were maintained on a bacterial lawn (*E. coli* OP50) on nematode growth medium (NGM) and incubated at 20^◦^C and 22^◦^C (details are provided in Table I) to ensure a continuous food supply. Worms were well-fed throughout the maintenance period as well as at the time of image and video acquisition, and no starvation was imposed at any point. Eggs observed on the plates were laid naturally by gravid adult worms maintained under the same culture conditions on NGM plates seeded with *E. coli* OP50, and no synchronization or external induction of egg-laying was performed. Experiments were performed with different population densities, which consisted of single as well as multiple worms (up to 20 worms approx) on the plate. Images were captured from different regions of the plate, which naturally leads to a variation in the number of worms per field of view. No external stimuli were applied to the worms during maintenance or at the time of imaging. A small fraction of mutant worms was used to add structural variations to the dataset. These were used only for model training and evaluation purposes, whereas the downstream behavioural analyses were performed using N2 worms only. The following mutants have been used in our dataset: FX05159 [*dma-1(tm5159)I*], NBR1170 [*shrIs3-Pmec4:chrimson:sl2:mcherry, goeIs5-pnmr1:GCAMP3.35*], and IK587 [*gcy-23(nj37) gcy8(oy44) gcy-18(nj38)*]. The first two, *i.e*., FX05159 and NBR1170, were used for training, and IK587 was used for prediction.

**Table I:** Dataset split across different experimental setups. The dataset consists of a total of 3018 images, comprising standalone images and frames extracted from a total of 114 videos, which we split across the training (2461 images), validation (400 images), and test (157 images) sets. The splitting was performed at the video level to prevent data leakage among these subsets. Exp 11 corresponds to the set of standalone images which were captured at multiple magnifications (2*×*, 3× and 4×) and varying backgrounds. The dataset has been provided in the [Google Drive](https://drive.google.com/drive/folders/1gVun2xOUyv7Z3EpdKyaNsZATIH460ExN?usp=sharing), and corresponding videos in these splits are specified in this [spreadsheet](https://docs.google.com/spreadsheets/d/1nIJoPMGl-PqyCVj39jKdRkCR5autmXSttdw9TKIjPfc/edit?usp=sharing).

| Experiments | Microscope | Temp. | Mutant/N2 | Train | Val. | Test | Total |
| --- | --- | --- | --- | --- | --- | --- | --- |
| Exp 1 | Leica M165 FC | 20 °C | N2 | 7 | 1 | 1 | 9 |
| Exp 2 | Leica M165 FC | 20 °C | FX05159 | 7 | 1 | 1 | 9 |
| Exp 3 | Leica M165 FC | 20 °C | NBR1170 | 13 | 3 | 2 | 18 |
| Exp 4 | Leica S9i | 22 °C | N2 | 28 | 5 | 3 | 36 |
| Exp 5 | Leica S9i | 22 °C | N2 | 3 | 1 | 1 | 5 |
| Exp 6 | Leica S9i | 22 °C | N2 | 7 | 2 | 1 | 10 |
| Exp 7 | Leica S9i | 22 °C | N2 | 2 | – | – | 2 |
| Exp 8 | Leica S9i | 22 °C | N2 | 2 | 1 | – | 3 |
| Exp 9 | Leica S9i | 22 °C | N2 | 2 | 1 | 1 | 4 |
| Exp 10 | Leica S9i | 22 °C | N2 | 16 | 3 | 1 | 20 |
| Exp 11 (images) | Leica S9i | 22 °C | N2 | 208 | 42 | 13 | 263 |
| **Images** |  |  |  | 2461 | 400 | 157 | 3018 |
| **Instances** |  |  |  | 5060 | 999 | 418 | 6477 |

A separate test dataset is prepared (we will call it “Test Dataset 2” wherever it is used), which contains videos that are slightly different from the training data. The dataset primarily contains adult N2 worms, with an experimental setup similar to that described in the previous paragraph. However, a small variability was added to test model performance in a bit challenging conditions, different from those used for training. For example, the test data contains videos of varying worm density, and the worms frequently collide and overlap with each other. Although the model is trained on overlapping worms, that fraction is very small compared to the total dataset. Therefore, the model appears to give better performance on single or isolated worms. We have also recorded videos with worms at different developmental stages, whereas the model has been trained strictly with adult worm images. In addition, we have also added various other kinds of variability, which include coloured patches on the background and different magnifications. Some mutant worms were also used (IK 587), which have a different postural morphology than N2. This dataset is used for model testing, which involves pose detection and evaluation of tracking accuracy, and not for annotation and training of the model. The corresponding datasets are provided and mentioned appropriately wherever they have been used. However, it should be mentioned that if the test images are significantly different in terms of resolution, intensity, contrast, *etc*, the model may lead to poor performance and model retraining may be required.

### B. Video and image acquisition

We acquired high-resolution videos of *C. elegans* using a Leica S9i stereomicroscope, equipped with an in-built CMOS camera, and a Leica M165 FC stereomicroscope with a Leica MC120 camera. These videos were subsequently used for model training and behavioural analyses. Imaging was performed at 2× magnification in both setups to ensure sufficient resolution for accurate pose detection and head-tail identification. The dataset consists of video recordings with both single and multiple worms under varying imaging conditions, as well as a range of simple and complex body postures. There are some standalone images (not extracted from any video) captured at different magnifications (2×, 3×, and 4×), and with varying backgrounds, to add variability in the training dataset. The details are provided in Table I.

### C. Annotations and dataset preparation

To train a deep learning model, we need to provide annotated images as input to the neural network. We extracted frames from videos of single and multiple worms, and manually labelled them in Roboflow^2^. Each worm was annotated by drawing bounding boxes around them, and a sequence of 11 keypoints were used to represent the body curvature. The key-points were placed along the approximate centerline of the worm body, starting from the head (kpt-0) and ending at the tail (kpt-10). Intermediate keypoints were distributed along the body in sequential order to capture the overall posture of the worm. We maintained this sequence, so that the model could learn about spatial distributions, patterns and ordering from raw images, which is essential for head and tail identification during pose estimation. A sample example of keypoints annotations is shown in Fig. 1 (top row, image on the right). The number of keypoints is chosen in such a way that it captures sufficient body-shape information, while minimizing annotation and training complexity. 11 keypoints are sufficient to capture common postures, including simple sinusoidal shapes as well as more complex bends such as omega and delta turns. Training with additional keypoints is computationally more expensive and may require better hardware for training. In the labelled dataset, each keypoint stores (*x, y, v*), *i.e*., the positional coordinates (*x, y*), and the visibility flag *v*, which can be set to 0, 1 or 2 depending on the situation. For example, when worms are partially out of the image, the out-of-frame keypoints are set to *v* = 0 by manually deleting those keypoints. When keypoints are within the frame, but are occluded by themselves or other worms, we set them to *v* = 1 by setting the flag to ‘invisible’. Keypoints that are properly visible and are within the frame (which is the most ideal case) are set to *v* = 2. During training and evaluation, the YOLO frameworks use the keypoints with *v* = 1 and 2 (giving equal importance to each, and not imposing any penalty to the occluded keypoints), while ignoring the invisible keypoints (*v* = 0). However, annotating the invisible keypoints is important as the model can learn real situations where worms can move out of the frame.

We annotated a worm only if at least 4 keypoints could be annotated. A total of 3018 images of single and multiple worms were annotated, which resulted in 6477 annotations. Out of the total dataset, approximately 82% images were used for training, and the remaining 13% and 5% were used for validation and testing, respectively. A summary is provided in Table I. Different data augmentation techniques, such as resize, auto-orient, grayscale, blur and noise were applied (check Table II for details) on the training set. This exercise helps in increasing the volume of training data as well as in adding variability to the dataset in order to enhance the model performance.

**Table II:** Details of the training process, including software versions, hyperparameters, and data augmentation methods. Augmentations were performed in Roboflow. Other hyperparameters were kept fixed at the default YOLOv8 values^a^ during training.

| Parameters | Values |
| --- | --- |
| Python | 3.12.7 |
| CUDA | 12.9 |
| Ultralytics | 8.4.3 |
| Model | YOLOv8m-pose.pt, YOLOv8l-pose.pt, YOLOv8x-pose.pt |
| Image size | $1024 \times 1024$ |
| Epoch | 1000 (with early stopping) |
| Patience | 100 (default) |
| Batch size | 18 to 36 (depending on the model and image size) |
| Optimizer | SGD (stochastic gradient descent) |
| Learning rate | 0.01 (default) |
| Momentum | 0.937 (default) |
| Weight decay | 0.0005 (default) |
| Seed | 0 (default) |
| Augmentation | $3 \times$ output<br>*Grayscale*: applied to 15% of images<br>*Brightness*: between $-15\%$ and $+15\%$<br>*Exposure*: between $-10\%$ and $+10\%$<br>*Blur*: up to 2.5px<br>*Noise*: up to 0.1% of pixels |

<sup>a</sup>https://docs.ultralytics.com/modes/train

### D. Dataset splitting

After annotations, we split the dataset before proceeding with training. To ensure a leakage-free evaluation, splitting was performed at the video level, so that the training, validation and test sets contain frames from different videos. The dataset classification is summarized in Table I, where videos corresponding to different experimental conditions are distributed among different splits. There were some standalone images (not extracted from any videos), recorded with Leica S9i microscope, which were also split among these subsets. As per our data split, 82% of the entire dataset was used for training, and the remaining 13% and 5% were used for validation and test, respectively. The entire training dataset has been provided in this [Google Drive](https://drive.google.com/drive/folders/1gVun2xOUyv7Z3EpdKyaNsZATIH460ExN?usp=sharing).

### E. Training and evaluation

We trained the YOLOv8 models for pose estimation using the annotated images. The training was performed on the training images, while the model performance was monitored on the validation set in each epoch. The final evaluation was performed on the test dataset to evaluate its generalization against ground truth annotations. To obtain the optimally performing model, we trained different YOLOv8 architectures for pose estimation, such as `YOLOv8m-pose.pt` (184 layers, 26.5M trainable parameters), `YOLOv8l-pose.pt` (224 layers, 44.5M trainable parameters) and `YOLOv8x-pose.pt` (224 layers, 69.5M trainable parameters) [69]. Different batch sizes were used based on the model architecture, image size, and available computational resources. All the training was performed on a fixed image resolution of 1024 × 1024. Other hyperparameters (such as optimization technique, learning rate, *etc*.) were kept fixed at the default values (details are provided in Table II). The training was performed on local hardware with Ubuntu 22.04 operating system and the following configurations: AMD Ryzen Threadripper 7970X processor, 128 GB DDR5 memory and dual NVIDIA RTX 4000 ADA 20 GB graphics unit.

Performance metrics corresponding to different YOLOv8 architectures on the validation set are mentioned in Table III. Standard metrics for object detections, such as precision (P), recall (R), F1 score, and mean average-precision (mAP), were used to evaluate performance. These metrics are calculated by comparing the model predictions with respect to the ground truth (GT) annotations, and keeping track of how many predictions actually match with GT (true positives), how many GTs are missed (false negatives), and how many objects are falsely detected (false positives). Here is a breakdown of each of these parameters.

**Table III:** Evaluation metrics for bounding box and pose detection for different YOLO models with 1024 × 1024 image sizes. Metrics corresponding to the highest mAP@0.5:0.95 for pose detection are shown.

| YOLO<br>architecture<br>(.pt) | Batch<br>size | Epoch | Bounding box metrics |  |  |  | Pose metrics |  |  |  | Training<br>time/epoch<br>(seconds) |
| --- | --- | --- | --- | --- | --- | --- | --- | --- | --- | --- | --- |
|  |  |  | P | R | mAP<br>@0.5 | mAP<br>@0.5:0.95 | P | R | mAP<br>@0.5 | mAP<br>@0.5:0.95 |  |
| yolov8m-pose | 36 | 901 | 0.989 | 0.993 | 0.995 | 0.956 | 0.958 | 0.962 | 0.974 | 0.837 | 210.8 |
| yolov8l-pose | 22 | 541 | 0.99 | 0.993 | 0.995 | 0.956 | 0.972 | 0.975 | 0.979 | 0.845 | 318.94 |
| yolov8x-pose | 18 | 276 | 0.989 | 0.996 | 0.995 | 0.949 | 0.969 | 0.965 | 0.97 | 0.816 | 468.18 |

- **Precision (P)** – It determines how many objects (worms, in our case) are correctly identified. It varies between 0 and 1 and is calculated as

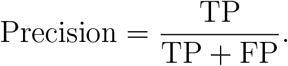

TP (True Positive) refers to predictions that actually match the GT objects. FP (False Positive) are those which the model predicts, but are not present in the GT. Higher P represents better prediction.
- **Recall (R)** – This metric takes into account how many true objects (present in the GT) are missed by the model. This metric also varies between 0 and 1 and is defined as

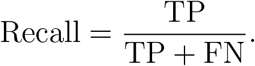

FN (False Negative) is the number of GT objects the model could not predict. The higher the value of recall, the fewer objects are missed by the model (lower FN), and hence the higher the detection accuracy.
- **F1 score** – This is a metric that combines precision and recall, and serves as a single-parameter performance metric of a model. This is calculated as the harmonic mean of precision and recall, which is given by

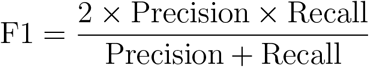
- **mAP@0.5** – Precision, recall and F1 score primarily deal with whether an object is correctly identified by the model. However, it is also important to determine the localization precision of the predicted objects. This is commonly measured using the mean average precision (mAP) metric, which evaluates the overlap between the predicted and ground truth bounding boxes. The overlap is measured by the intersection over union (IoU), defined as the ratio of the area of overlap to the area of the union between the predicted and GT bounding boxes. In particular, YOLO models return mAP@0.5, where a detection is considered a true positive only when the IoU with the corresponding GT bounding box is greater than or equal to 0.5. The metric is defined as

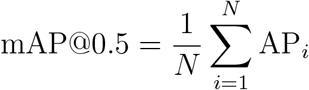

where AP_*i*_ denotes the average precision for the *i*^th^ class, calculated from the area under the precision-recall curve for detections that satisfy the above condition. *N* is the total number of classes in the dataset. A higher mAP@0.5 value indicates better detection and localization precision.
- **mAP@0.5:0.95** – Although mAP@0.5 is a useful measure of detection performance, evaluating performance at a single IoU threshold may not fully capture localization precision. To address this limitation, we additionally report mean average precision at multiple IoU thresholds, commonly termed as mAP@0.5:0.95. This metric evaluates average mAP at 10 different IoU thresholds, ranging from 0.5 to 0.95, with a step size of 0.05. This is defined as

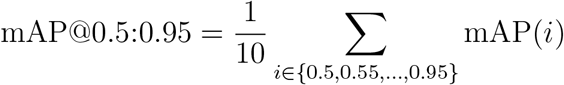

Since this metric evaluates the prediction at multiple thresholds, it provides a more comprehensive assessment of detection and localization precision.

### F. Handling collisions and overlaps

During pose detection and tracking of multiple worms, scenarios may occur where worms collide with each other, causing overlaps and occlusions. An overlap is defined as the event in which two or more worms come so close to each other that their body segments become inseparable, and the associated keypoints indistinguishable. During tracking, the worm identities are maintained across frames by using the in-built “ByteTrack” or “Bot-SORT” tracking algorithms. They maintain worm identities across consecutive frames by first predicting the worm locations in the next frame using a Kalman filter. The predicted bounding boxes are then compared with the newly detected boxes using the IoU metric. Finally, the Hungarian assignment algorithm [70] is applied to optimally associate detections with predictions, thereby preserving consistent identities over time. The model can preserve the identities when worms partially or moderately overlap with each other; however, for heavily overlapping situations, particularly when we track worms in crowded environments, it becomes challenging to maintain those identities, resulting in identity switch and continuity loss. We have reported tracking metrics such as MOTA, MOTP and IDF1 scores for worms at different populations to demonstrate the model performance and the extent to which it produces reliable outputs.

## III. RESULTS

Now we present the key results of this study. This section is divided into two parts. In Subsec. III A, we describe the performance of the DPT model on the validation and test datasets. In addition, we report the tracking metrics on test videos with varying experimental conditions. In the next Subsec. III B, we demonstrate how DPT has been integrated with several behavioural analyses, viz., pose detection, tracking, motion orientation, identification of forward motion and reversals, detection of omega turns and finally, detection of multi-class objects (eggs and worms) within images.

### A. Model performance and accuracy

During training, YOLO automatically generates two different weight files – `best.pt` and `last.pt`. As the names suggest, the `best.pt` corresponds to the model checkpoints that contributed to the best validation performance at any epoch during training. Similarly, the `last.pt` corresponds to the checkpoints at the last epoch. The model performance is determined based on the fitness score, which is calculated as

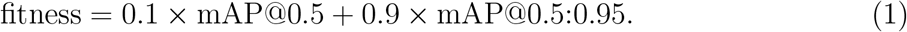

The best-performing model corresponds to the checkpoints with the highest fitness score, which is primarily determined by the mAP@0.5:0.95 metric. YOLO pose models predict bounding boxes and keypoints, and hence metrics for both are estimated. But, for pose detection tasks, the pose metrics are given priority. The model evaluations for bounding box and keypoints predictions are given in Table III, corresponding to the highest mAP@0.5:0.95 metric for pose detection. For all follow-up behavioural analyses and quantification, we used the `best.pt` model, which we renamed later according to the model name and image size.

Different YOLOv8 pose architectures were trained to obtain the best-performing model on our custom dataset. We used high-resolution images for training. Training with low-resolution images can be faster, but it risks losing important spatial features, which can lead to poor performance. Similarly, a larger model can learn patterns in the data more effectively than a smaller one. The training was initialized with 1000 epochs; however, the process was automatically terminated at some lower number of epochs using early stopping, as reported in Table III. The training process stops when the model does not show any improvement over a certain number of epochs on the validation dataset. This helps reduce computation time and also prevents overfitting. According to the table, precision and recall for bounding box prediction are ~0.99, with mAP@0.5 ~0.995 for all models, indicating that worms are correctly detected with very high localization precision. The pose detection task focuses on keypoints detection along the body of *C. elegans*, which is more complex than object detection. The precision and recall for the pose detection vary between 0.96 and 0.975, with mAP@0.5 ranging between 0.97 and 0.98, which is significantly high and appropriate for behavioural quantification like eigenworms decomposition, reversal and omega turns detection, *etc*. The metric mAP@0.5:0.95 corresponding to the best model is ~0.95 for bounding box prediction. For pose detection, it varies between 0.816 and 0.845. This is lower compared to mAP@0.5, as we consider multiple IoU thresholds for evaluation, which we discussed earlier. However, pose detection models with mAP@0.5:0.95 value higher than 0.8 are suitable for deployment and applications, as is also evident for pretrained YOLO pose models on the COCO-keypoints dataset^3^. The accuracy of pose detection as reported in the table refers to the standard detection metrics on the evaluation dataset, and the accuracy of predicted keypoints locations relative to the annotated ground truths. In addition to precision, recall and mAP values, we have also measured the performance on the test dataset and videos. Corresponding inference times and frame rates for each of the models are summarized in Table IV. It should be noted that the reported metrics are evaluated on the validation and test datasets. The model can show poor performance if tested on images that are significantly different from the training dataset used in this study.

**Table IV:** Inference time and frame rates of different YOLO models on test data (with batch size of 16) and videos containing single as well as multiple worms. Model names are changed according to the network architecture and image size.

| Model | Image | Test data (Images) |  | Video 1 (1 worm) |  | Video 2 (9 worms) |  |
| --- | --- | --- | --- | --- | --- | --- | --- |
|  |  | Inference time (ms) | FPS | Inference time (ms) | FPS | Inference time (ms) | FPS |
| yolov8m-pose-1024.pt | 1024 | 25.9 | 38 | 10.7 | 93 | 10.7 | 93 |
| yolov8l-pose-1024.pt | 1024 | 41.5 | 24 | 17.3 | 57 | 17.3 | 57 |
| yolov8x-pose-1024.pt | 1024 | 61.8 | 16 | 25.9 | 38 | 26.2 | 38 |

In addition to showing the best-performing metrics, the performance is also monitored in each epoch, and the results are shown in Fig. 2. Here, we have shown the pose metrics only, since we are primarily interested in keypoints detection. The precision, recall and mAP@0.5 grow rapidly, and reach ≥0.9 within the first 50 epochs, and ≥0.95 within 150 epochs, showing strong performance and rapid convergence of the accuracy. In the remaining epochs, the accuracy gradually increased to the maximum values. The metric mAP@0.5:0.95 takes relatively longer to converge, as shown in Fig. 2D. The best performance is obtained with `yolov8l-pose.pt` model, according to the pose metrics reported in Table III. Therefore, we use this model for all follow-up analyses and quantification, except for omega turns detection. The model `yolov8l-pose.pt` has overall better accuracy during training; however, `yolov8x-pose.pt` was observed to give better predictions of complex postures such as omega turns. There is a tradeoff between prediction accuracy and inference speed for each model, as smaller models enable faster inference, but may come at the cost of reduced accuracy. It should also be noted that all the metrics and reported results are evaluated within the validation and test sets, which contain images of similar experimental conditions as described in Table I.

**Figure 2:**
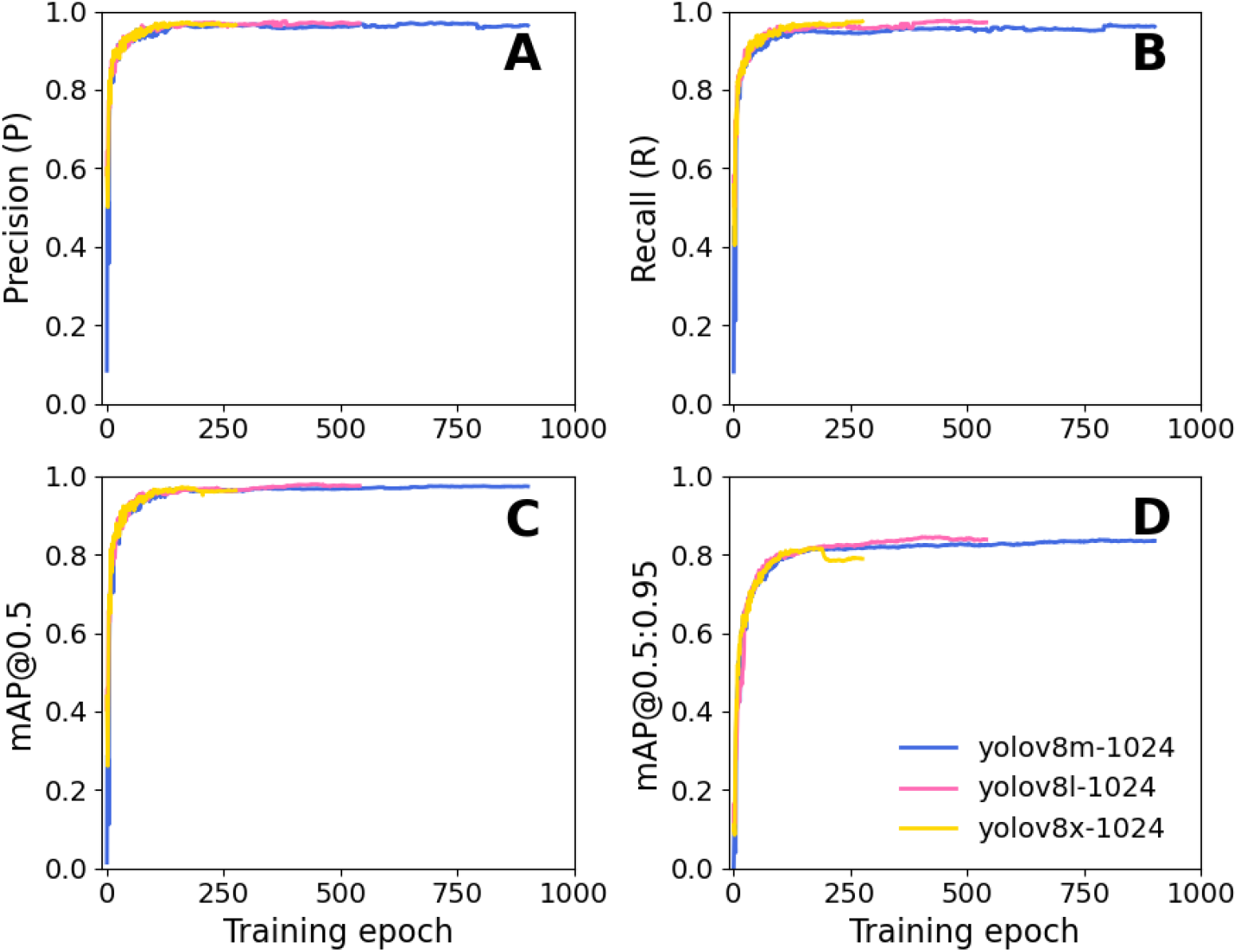
Model performance on validation dataset during training. Pose metrics for keypoints detections of *C. elegans* with different YOLOv8-pose architectures are shown. Corresponding evaluation metrics, such as precision, recall, mAP@0.5, and mAP@0.5:0.95 at different training epochs are shown.

#### 1. Tracking accuracy

Tracking objects accurately is one of the most essential yet challenging task of any deep learning model. During object tracking, unique identities are assigned to the objects, and those identities should remain consistent throughout the video for accurate tracking. But, inconsistencies may appear when objects overlap with each other, or an object loses identity because of very low visibility or frame rates. To evaluate the tracking accuracy, we have measured standard tracking metrics like MOTA (Multiple Object Tracking Accuracy), MOTP (Multiple Object Tracking Precision) and IDF1 (Identification F1 score) on different videos with varying worm populations (definitions are described later). The metrics are evaluated by comparing the predictions with annotated ground truths (GTs), which were prepared by manually drawing bounding boxes around each worm in every frame of the video using the web-based Computer Vision Annotation Tool (CVAT)^4^. During annotations, every object is associated with a unique tracking identity, and the identity is maintained in each frame throughout the video. Those identities and associated bounding box properties are then used as references to measure various tracking metrics as mentioned above. Only adult worms were annotated to remain consistent with the keypoints annotation framework.

A total of 7 videos from Test Dataset 2 (as described in Sec. II A) with different populations and varied overlap scenarios were used to evaluate tracking performance, and the summary is provided in Table V. We used `yolov8l-pose-1024.pt` model for tracking, and accuracy was evaluated using the motmetrics^5^ library following the standard MOTChallenge evaluation protocol [71, 72]. In each frame, a pairwise cost function is measured between the predicted and ground truth bounding boxes to find one-to-one matching between each object by calculating the intersection-over-union (IoU) metric [73]. Specifically, a cost between the predicted (*i*) and GT (*j*) boxes *C*_*ij*_ = 1 −IoU_*ij*_ is measured, and an IoU threshold (0.5 in our evaluation) is used, so that any pair with IoU below the threshold is considered invalid and excluded from evaluation. This cost matrix is then used to compute one-to-one association between the predicted and ground truth objects using the Hungarian algorithm on each frame [70]. These associations are then used to identify whether a detection is considered a true positive, true negative or false positive. Additionally, consistency between predicted and GT identities are also checked, and it keeps track of any identity switch that happens over time. These frame-wise associations are collected across all frames to calculate the standard tracking metrics like MOTA, MOTP and IDF1 using the following formulas:

**Table V:** Tracking metrics such as MOTA, MOTP and IDF1 measured on videos containing different numbers of worms. We used the `yolov8l-pose-1024.pt` model for tracking, and metrics were measured with an IoU threshold of 0.5. Reference videos are shared here.

| Video file | #worms | Inference time | FPS | MOTA (%) | MOTP (%) | IDF1 (%) |
| --- | --- | --- | --- | --- | --- | --- |
| video1.mp4 | 1 | 16.1 ms | 62 | 100 | 92.7 | 100 |
| video2.mp4 | 2 | 15.7 ms | 63 | 100 | 93.4 | 100 |
| video3.mp4 | 2 | 15.5 ms | 64 | 62.7 | 86.7 | 62.7 |
| video4.mp4 | 2–5 | 15.7 ms | 63 | 86.1 | 91.5 | 92.9 |
| video5.mp4 | 3–6 | 15.6 ms | 64 | 80.7 | 90.2 | 90.4 |
| video6.mp4 | 4–9 | 15.4 ms | 64 | 90.5 | 91.8 | 94.9 |
| video7.mp4 | 14–17 | 15.1 ms | 66 | 85.9 | 92.4 | 88.8 |

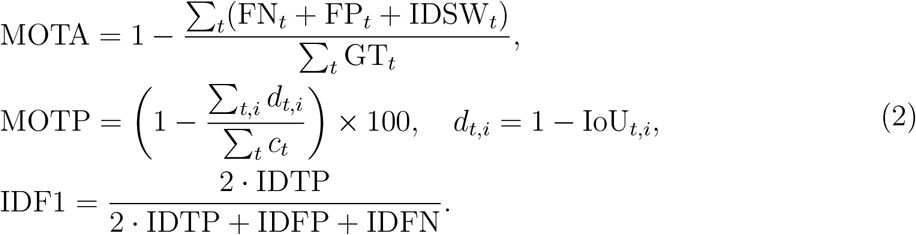

MOTA is calculated by finding all false negatives (FN), false positives (FP), identity switches (IDSW), and the number of ground truth (GT) objects, summing over all frame sequences *t*, and subtracting from 1. MOTP corresponds to the localization precision of the matched pairs, computed from the overlap between the predicted and GT bounding boxes. The tracking accuracies were evaluated using the motmetrics library, which computes MOTP as the average localization error, calculated from the localization distance defined as *d*_*t,i*_ = 1−IoU_*t,i*_ and averages over all matched pairs across all frames [71]. However, in this work, we report MOTP as localization accuracy, calculated using the modified formula mentioned in Eq. (2), to express the metric as a percentage-like value for easier comparison with MOTChallenge-style evaluations [72]. Therefore, the reported MOTP corresponds to the average overlap (IoU) between the predicted and GT bounding boxes, with higher values indicating better accuracy. IDF1 shows the model’s consistency in maintaining the same identity. It is calculated by keeping track of all the identities (ID) given to the TPs, FPs, and FNs.

Table V consists of videos with a minimum and maximum number of worms present at any instant of the video. The evaluation code and dataset are provided on [GitHub](https://drive.google.com/drive/folders/1es_7pOL9x9ipI3vDHsXvjmSvo8zcaouS?usp=sharing). For a very small population, *e.g*., in video1 and video2, no overlap happens, which results in high tracking accuracy. But, for video3, the worms heavily collide and overlap with each other, causing frequent identity loss, which results in very low accuracy. Similarly, in video4 and video5, worms moderately overlap with one another, resulting in moderate accuracy scores. The reason is that the model sometimes detects the small worms (probably in the L1 or L2 stage), which we did not annotate during the GT preparation, and those detections are considered false positives, causing low precision. Similarly, in relatively large populations, *e.g*., in video6 and video7, worms undergo collisions, which reduces the accuracy score. But there are other worms which do not encounter any collision, and no identity switch happens.

It helps maintain an overall good accuracy score. Therefore, the model achieves good overall tracking performance on test videos, maintaining reliable multi-worm tracking even in the presence of moderate collisions and overlaps.

In crowded environments, where worms frequently collide and overlap, it becomes difficult to track them accurately. However, several instances have been observed where the algorithm is able to maintain the tracking identity. For example, in video3 in [Google Drive](https://drive.google.com/drive/folders/1rtKqQcFahmdZ3F8tpkPazOQaBAp_8qFc?usp=sharing), worms 1 and 2 collide with each other, but worm ID 1 is maintained during their first collision. However, in the second collision, the ID is lost. Similarly, in video 6, worms 6 and 8, and worms 4 and 7, collide with each other, but identities are maintained throughout the video, resulting in a high IDF1 score. However, in the case of images or videos that are significantly different from training images, the accuracy can degrade, and model retraining on the representative dataset would be required.

### B. Deep-Pose-Tracker (DPT)

Let us now discuss some key results of predicted worm postures and the development of DPT-based behavioural analysis tools for *C. elegans*. The quantities we report are as follows: (1) pose detection, (2) eigenworms decompositions, (3) worm tracking and speed measurement, (4) spatial exploration measurement, (5) orientation of motion, (6) forward-reverse locomotion, (7) quantification of omega turns, and (8) multi-class object detection and counting.

#### 1. Pose detection

Pose detection of model organisms like *C. elegans* are important in the context of behavioural cues for research in the areas of neuroscience, genetics and drug discovery, among others. It involves the detection and quantification of body shape variations of these worms from images or videos. Pose detection is useful in identifying the movement patterns of *C. elegans*, particularly in the context of roaming and dwelling behaviours [30, 74], in response to external stimuli [75] and understanding how they correlate to various molecular mechanisms [76, 77]. For example, in the roaming state (also known as global search), worms tend to travel long distances by moving fast while making small body bends. On the other hand, in the dwelling state (also known as local search), they continue to move in a localized region, hence move very slowly and make more body bends than in the other (roaming) case [74, 78]. They also occasionally exhibit reversal movements, which are often associated with complex body bends, such as omega and delta turns [79], which are again related to the posture dynamics of *C. elegans* as responses to external stimuli.

We used DPT model to perform pose detection tasks on single and multiple worm videos, which is the primary component of this work. Performance of the model is checked on the test dataset, which gave the following metrics: precision = 0.95, recall = 0.96, mAP@0.5 = 0.96 and mAP@0.5:0.95 = 0.82. Some sample images with predicted keypoints are shown in Fig. 3A. Keypoints are shown by coloured circular dots starting from the head (kpt-0), denoted by a blue dot at one end, to the tail (kpt-10), denoted by a green dot at the other end. The model is able to capture different body postures of *C. elegans*, which include simple sinusoidal shapes to complex bending like omega turns. However, the model occasionally fails to make accurate predictions for worms exhibiting highly coiled postures. The keypoints coordinates are stored, which are further used for several downstream analyses (discussed in the following subsections). The model can be efficiently used on images and videos and is also suitable for multi-worm pose detection. Some examples of pose detection on single and multi-worm videos from the test dataset are shown here. Model performance is checked on videos from the Test Dataset 2 as well, which are slightly different from the training images, as described earlier in Sec. II A. It should be noted that these videos are not used for training and therefore do not have any ground truth annotations. This ensures that performance is checked under conditions different from the ones on which it has been trained and tested earlier. Some representative images are shown in Fig. 3B. Here, predictions of worm postures under different experimental conditions, such as background noise, crowded populations and overlaps, different magnifications, mixed population containing adult and L3-L4 worms, and mutant worms (IK587), are shown. In all these cases, the model predicts body postures with reliable head-tail identification. There are several instances where worms touch or partially overlap with each other, for example worms 3-4 in Fig. 3A4, worms 2-4 in Fig. 3B2, worms 6-9, 14-17, and 12-15 in Fig. 3B3, but the model predicts their poses separately. However, accurate pose detection and tracking in severe occlusions are challenging and may require further training with images having more complex body postures and overlaps.

**Figure 3:**
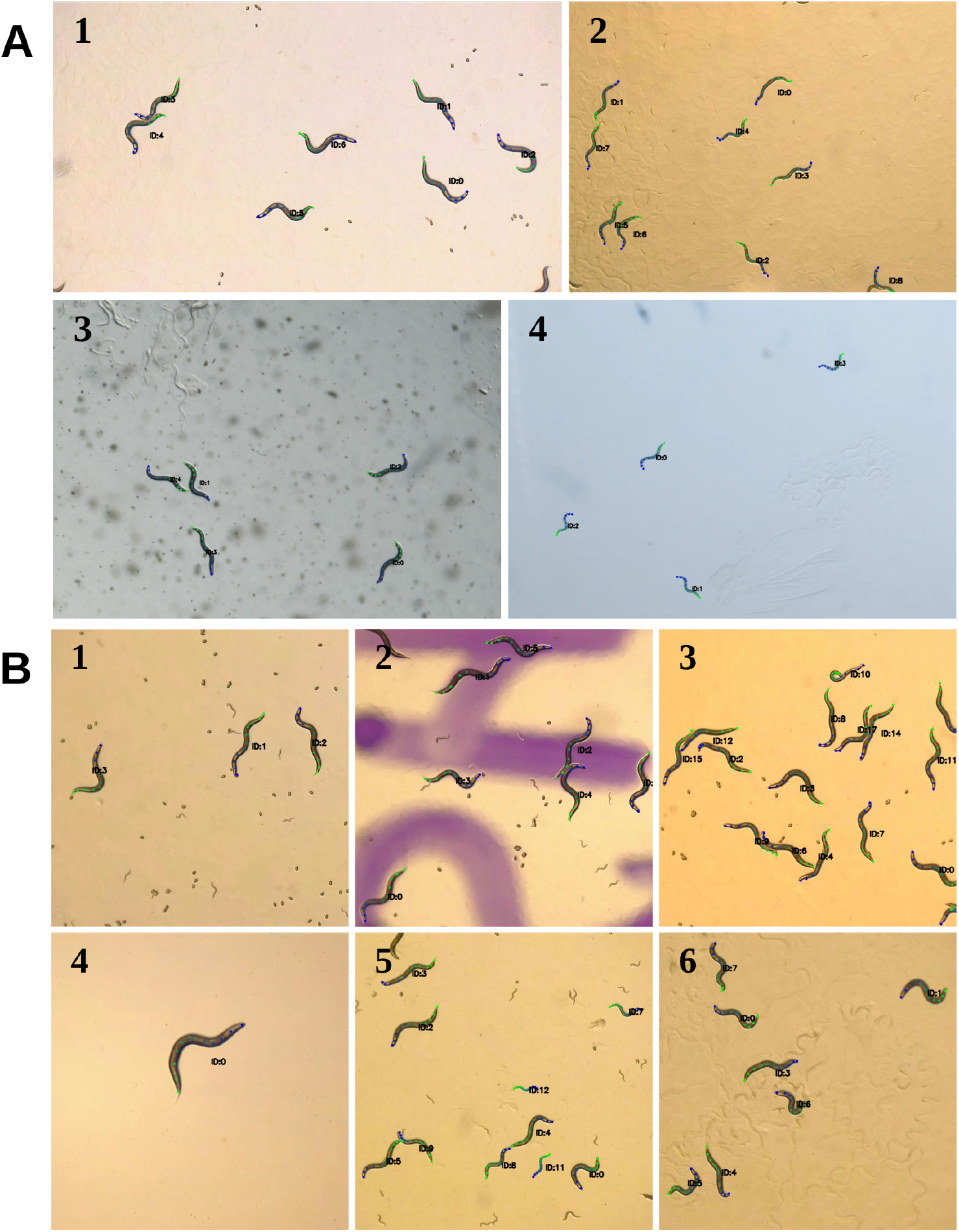
Sample images of pose detection of *C. elegans*. (A) Pose detection with varying backgrounds, image contrast, noise, and worm sizes, within the test dataset. Detected poses are shown with keypoints along the body centerlines, with the blue dot (kpt-0) at one end representing the head, and the green dot (kpt-10) at the other end representing the tail of the worms. (B) Model performance on Test Dataset 2. Pose detection on various imaging conditions, such as 1. in the presence of background noise (eggs and small worms), 2. patches in the background, 3. crowding and overlaps, 4. different magnification, 5. mixed population containing adult and L3-L4 worms, and 6. mutant worms (IK587). Sample examples of pose estimation on images and videos are available in the Supplementary Materials (S1 and S2) and in the [Google Drive](https://drive.google.com/drive/folders/15q9nzsPEaV9KlkzCwJFaTdMuFrQviGX3?usp=drive_link).

#### 2. Eigenworms decompositions

Quantification of *C. elegans* locomotion is tedious due to their complex movement pattern. To perform locomotion, the worms bend their body in a sinusoidal pattern, which creates a wave-like posture along the body. They occasionally make more complex body bends, like omega or delta turns, when they are exposed to unfavourable conditions in the surroundings.

Analytically capturing these pose variabilities by any brute force method would require a large number of degrees of freedom, which may not be effective in every situation. However, these complex locomotion patterns can be quantified more effectively using a few fundamental modes, known as *eigenworms* [21, 68, 80, 81]. The entire body shape variation and posture dynamics can be represented by these eigenworms, with the help of a few degrees of freedom.

As the DPT model is trained to predict worm postures, a straightforward extension is to evaluate eigenworms using the predicted body shapes. We used the keypoints data of day 1 adult *C. elegans* (N2) to quantify eigenworms from multiple videos. As discussed previously, we start by presenting a sample with its predicted pose in Fig. 4A, which contains 11 keypoints distributed along the body. Evaluating eigenworms is based on finding the correlation between the neighbouring body segments. The correlation can be evaluated by measuring the tangent angles between the segments, followed by constructing a correlation matrix and finding the most significant modes from the matrix. But with 11 keypoints, the tangent angles are highly discontinuous, and it is difficult to find any meaningful pattern. To overcome this issue, we increase the number of keypoints by using spline interpolation, from 11 to 101. The result is shown in Fig. 4B, in which red dots are the actual keypoints, and blue dots are interpolated points, which are made to be equidistant from each other, essential to get a smooth correlation matrix. In Fig. 4C, we have shown the tangent angles (with the horizontal axis) between every keypoint shown by a colour scheme, and the colourbar represents the magnitude of these angles. The tangent angles are calculated as *θ* = tan^−1^ (*dy/dx*), where *dx* and *dy* are the differences in coordinates between two subsequent keypoints. The head of the worm is indicated by the red dot. It should be noted from Fig. 4C that, starting from the head, the magnitudes of angles do not change randomly; rather, they gradually change from a high to a low value, then again to a high value. This smooth variation is present because of the fact that the dynamics of body segments are correlated during their motion, which in turn allows dimensionality reduction and describes the movement patterns in a low dimension. The posture of the worm must be reoriented so that the tangent angles do not depend on the orientation of the worm, and the curvature becomes an inherent property of the body shape only. This can be done by simply subtracting the mean value of *θ* from *θ* itself, and the result is shown in Fig. 4D. This reoriented *θ* is then used to evaluate the covariance matrix, and the principal components governing the eigenworms.

**Figure 4:**
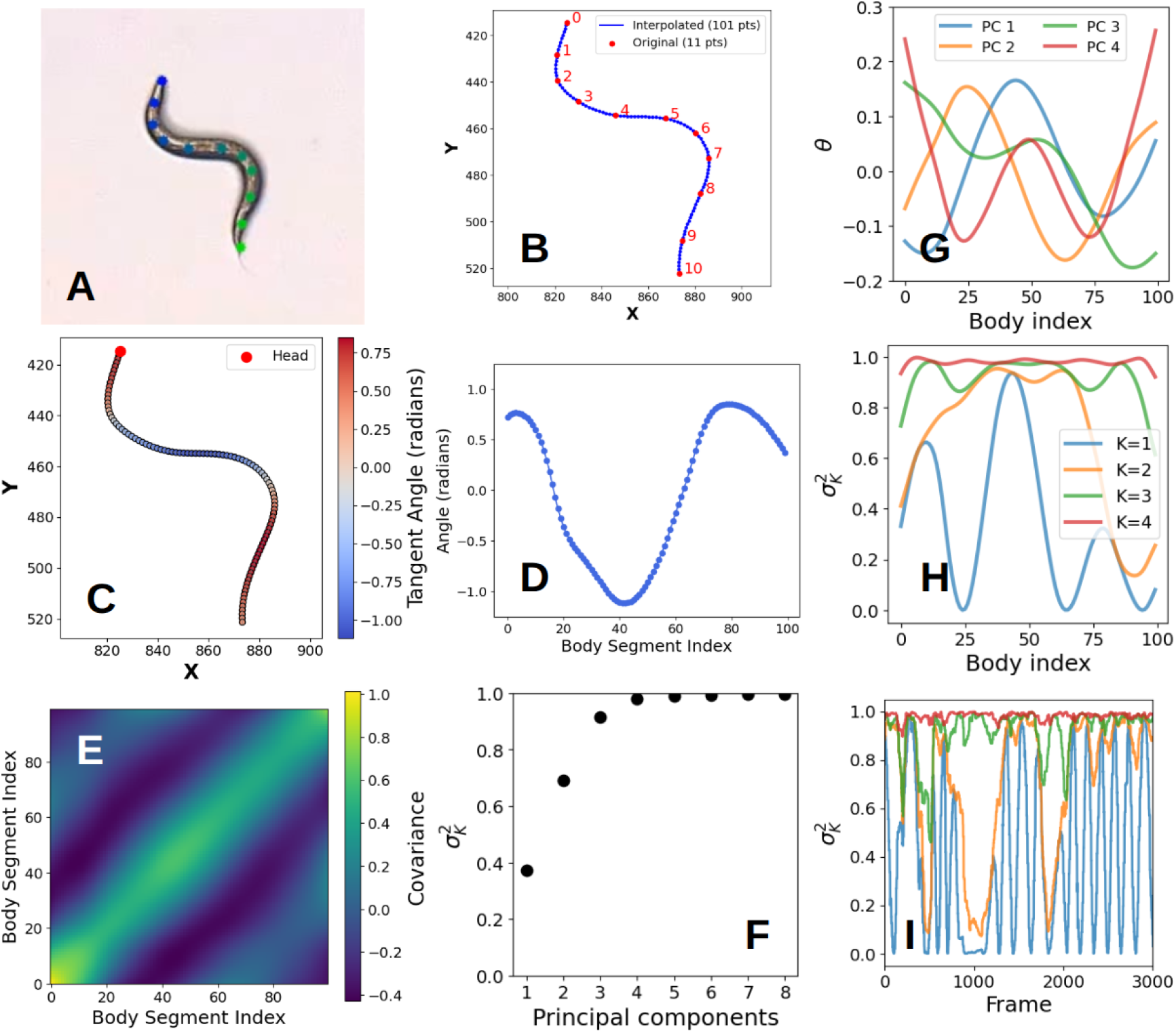
Quantification of eigenworms from body postures of *C. elegans*. (A) The 11 keypoints along the body of the worm. The blue and green dots at the ends represent the head and tail of the worm, respectively. (B) Spline interpolation is used to create 101 keypoints to get smooth characteristics of the tangent angles. (C) Tangent angles are represented by colours. The colour bar shows the magnitude of the tangent angles. Variation of colour along the body shows how the tangent angles of different body segments are correlated with each other. (D) Tangent angles are subtracted from their mean value to make them independent of the orientation of the worms. (E) Covariance matrix showing the correlation between different body segments. (F) The fraction of an eigenvalue over the sum of all eigenvalues. The first four modes capture more than 95% of the total variance. (G) Corresponding principal components (PCs) or eigenmodes, also known as the eigenworms. (H) The first 4 eigenworms are able to capture the geometrical variations of the worm. Variance captured with *K* eigenmodes, with *K* varying from 1 (bottom) to 4 (top) to show, for each of them combined, how much variance is captured compared to the total variance. (I) Dynamic variations of body curvatures are represented by the leading 4 eigenworms. Similar to Fig. H, this figure shows the combined contributions of the leading eigenworms (from bottom to top). The dataset has been provided in the [Google Drive](https://drive.google.com/drive/folders/1pMhO9ieUpDSa4jaKXjWMCiVqWlnr2EP1?usp=sharing).

Now we demonstrate the most important step for eigenworms decomposition and determining the most significant modes in the movement patterns. First, we construct the covariance matrix *C*(*s, s*′), using the orientation-independent tangent angles along the body of the worms. Here, *s* and *s*′ represent the body segments. *C*(*s, s*′) is a 100 × 100 symmetric matrix which is evaluated by measuring [68]

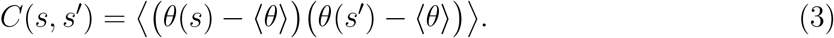

The covariance matrix, as shown in Fig. 4E, is computed from multiple videos of N2 worms, consisting of different types of body postures. It includes ordinary sinusoidal movements, forward-reverse motions, different types of body bends, head and tail movements, *etc*., which are required to obtain all different types of shape variations. A total of ~104k frames from multiple videos are used to construct this matrix. The covariance matrix shows a very strong correlation along the diagonal axis, which indicates that movements of all the body segments are correlated with their neighbouring segments. These patterns in the matrix indicate strong correlations of different body segments of *C. elegans* which are characterised by their movements.

The patterns in the covariance matrix *C*(*s, s*′) also indicate that a few of the modes have higher contribution towards the overall movement. An intuitive way of finding these modes is to calculate the eigenvalues of *C*(*s, s*′), and evaluate the fraction of a particular eigenvalue compared to their sum. This can be done as

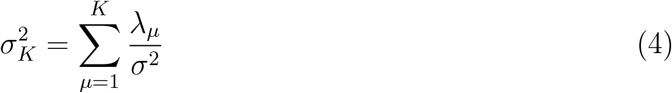

where 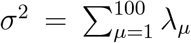, the sum of all eigenvalues, and *K* is the number of modes. This has been arranged in descending order and has been plotted in Fig. 4F. Here, *λ*_*µ*_ is the eigenvalue of *C*(*s, s*′) obtained from *C*(*s, s*′)*a*_*µ*_ = *λ*_*µ*_*a*_*µ*_, where *a*_*µ*_ represents the eigenvectors. From Fig. 4F, it is evident that 4 principal eigenmodes capture more than 95% of the total variance in movement patterns. It means that although the movement patterns are inherently complex, there are some correlations, which make it possible to describe complex locomotion in a low-dimensional space. In Fig. 4G, eigenvectors corresponding to each of the significant modes are shown, which are also known as the *eigenworms*. The first two eigenworms represent the sinusoidal motions that worms exhibit when they move. It appears as though two sinusoidal waves propagate along the body with a phase lag, as the worms move, and they contribute to almost 70% of the entire variance. The third one corresponds to body bendings, such as omega and delta turns. The fourth eigenworm represents the movement of the head and the tail regions.

Now, it should be checked whether these principal components can actually describe the overall geometric variations of the body and the posture dynamics. First, we reconstruct the worms’ body curvatures by combining 4 primary eigenmodes, which is shown in Fig. 4H. The first mode is a sinusoidal wave with varying amplitude along the body. As we combine more eigenmodes, it captures more variations in body shapes. For example, a combination of 2 eigenmodes can represent almost 70% of the total variance, as evident from Fig. 4F. When the third eigenmode is added, it captures the variabilities coming from the first and last 25 segments (approximately) of the body, and captures almost 90% variations. But the variations near the heads and tails are still missing, which are captured by combining the fourth eigenmode. Therefore, the combination of these eigenworms represents the geometric curvature along the body of the worm in any frame. It should also be checked if this combination is sufficient to describe the posture dynamics, *i.e*., the body shape variations over time. To demonstrate that, we have shown the combinations of these 4 eigenmodes as a function of frame number (which is equivalent to time) in Fig. 4I, which shows strong resemblance to the total variance.

#### 3. Worm tracking and speed measurement

Detection and tracking are among the most fundamental and useful tasks in studying behaviours of *C. elegans* or any other model organism. It requires quantitative analysis of videos to track individual worms under various experimental conditions. In any standard tracking process, the objects are assigned unique identities, which are maintained throughout the video. The process of identity assignment is already described in Subsubsec. III A 1. For tracking a single object, maintaining the ID is not difficult. However, in multi-object tracking systems, the assigned ID should remain consistent across frames. Developing a tracking model for *C. elegans* is difficult, as they are visually identical and often undergo collisions and overlaps, which makes it extremely difficult to maintain their identities. This challenge has been discussed previously, and tracking metrics with different worm populations are shown in Table V. In Deep-Worm-Tracker, Banerjee et al. integrated the StrongSORT algorithm with YOLOv5 to keep track of individual worms across frames [33]. The upgraded version, YOLOv8, has integrated tracking algorithms^6^ within it, eliminating the need for incorporating external tracking modules [39, 43, 47]. We used the `botsort.yaml` module, which assigns unique IDs to the detected objects and keeps track of them in each frame of the video. We provide videos of *C. elegans* as input to our tracker model, which returns the coordinates of each worm in every frame. From those coordinates, we obtain the trajectories of individual worms. A sample result of multi-worm trajectories from a single video is shown in Fig. 5A. Once we obtain the trajectories, we can evaluate several physical quantities, such as the distance travelled by a worm, locomotion speed, area explored during their movements, and orientation, which we discuss later. These quantities are important for understanding and interpreting the behavioural patterns of model organisms under various external conditions and internal perturbations. To track *C. elegans*, we used the trained DPT model, as it performs object detection and tracking in addition to pose estimation. But, if we are interested only in tracking and related behaviours, then the object detection models of YOLO (instead of keypoint detection) are more appropriate, as they will be less computationally expensive.

**Figure 5:**
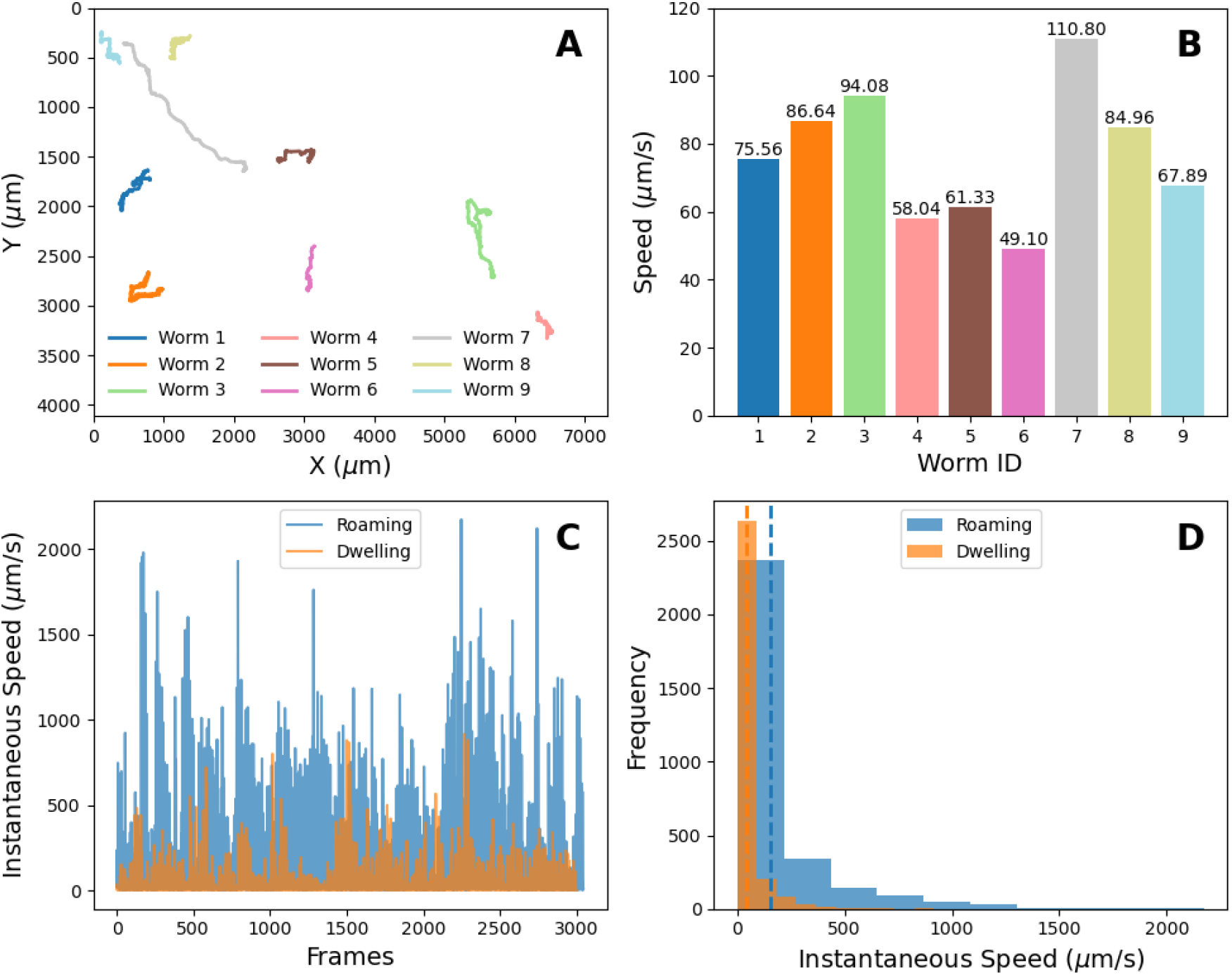
Measuring speed from a video containing multiple worms. (A) Trajectories of individual worms and (B) corresponding movement speed for each trajectory, shown with the same colours. (C) Instantaneous speed of worms’ locomotion at each frame corresponding to the roaming and dwelling behaviours. (D) Distributions of instantaneous speeds for roaming and dwelling movements. The vertical dashed lines indicate the mean values: 40 *µ*m/s for dwelling, and 155 *µ*m/s for roaming behaviours. Relevant data are shared in the [Google Drive](https://drive.google.com/drive/folders/1gw1ochBa90ZzvIvy88fvHeW-wLl6Scvk?usp=sharing).

As an integral part of tracking, we have also measured the locomotion speed of worms from their trajectories, as this is one of the most important features in behavioural studies. We calculate the distance travelled by worms in each frame using *dx_i_* = *x_i_* − *x_i_* − 1 and *dy*_*i*_ = *y*_*i*_ − *y*_*i*−1_. The displacement of the worm between two consecutive frames is then calculated using 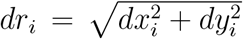, and added to give 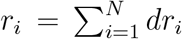, the total distance travelled by the worms. This total distance is used for calculating the average speed using *v* = *r/t*, where *t* is the total duration of the video. The distance and speed are measured in standard units (*µ*m) by taking the suitable scaling parameter for the microscope and camera module that has been used for data acquisition. A resulting output of locomotion speeds is shown in Fig. 5B. Here, the worm IDs are the same as the trajectories shown in Fig. 5A. Similarly, we can measure the instantaneous speed to show framewise movement of the worms, which is shown in Fig. 5C, and the associated distributions are shown in Fig. 5D. These quantities are often useful to distinguish roaming and dwelling behaviours. The corresponding videos are provided in the figure caption.

#### 4. Spatial exploration measurement

In addition to measuring distance and speed, it is also important to determine the extent of spatial exploration the worms carry out during their motion. However, given that we are already measuring the distance, what additional information does this spatial exploration (SE) measurement provide? In some cases, SE can proportionally change with distance, but in the context of global and local search, also referred to as roaming and dwelling movements, this intuition can break [30, 82, 83]. During local search, worms tend to travel less, and their locomotion is confined to a small region. Although the locomotion happens in a confined space, their bodies keep moving within that region, resulting in covering a large distance. On the other hand, in global search scenarios, worms travel long distances in search of food, hence exploring a larger space. Besides these two, other situations may appear when a worm moves in a loop-like trajectory and returns close to the initial position, resulting in travelling a large distance but exploring a relatively smaller space. In such a situation, measurement of spatial exploration, along with distance, provides a more accurate description of global and local search behaviours.

Several methods can be utilized to calculate SE by the worms during their motion. The simplest method (Method 1) is to draw an imaginary rectangle around the trajectory of the worm. Then, calculate the area of that rectangle as shown in Fig. 6A, which would represent the region explored by the worm. However, this method can sometimes lead to incorrect measurements, as it is biased towards the shape and orientation of the trajectory. Imagine a situation where a worm travels a long distance, but the trajectory is approximately horizontal or vertical. It would have a very small height or width, resulting in nearly zero area measured by this method. This is certainly not true, as the measurement can be severely underestimated sometimes. Therefore, an alternate approach should be used, which is independent of the orientation and shape of the trajectory on the XY plane. We, therefore, design Method 2, which calculates a distance measure reflective of the spatial exploration. To do this, we first calculate the projections of the trajectory, *R*_*x*_ and *R*_*y*_, on each of the axes, as shown in Fig. 6B. Then we can calculate SE as

**Figure 6:**
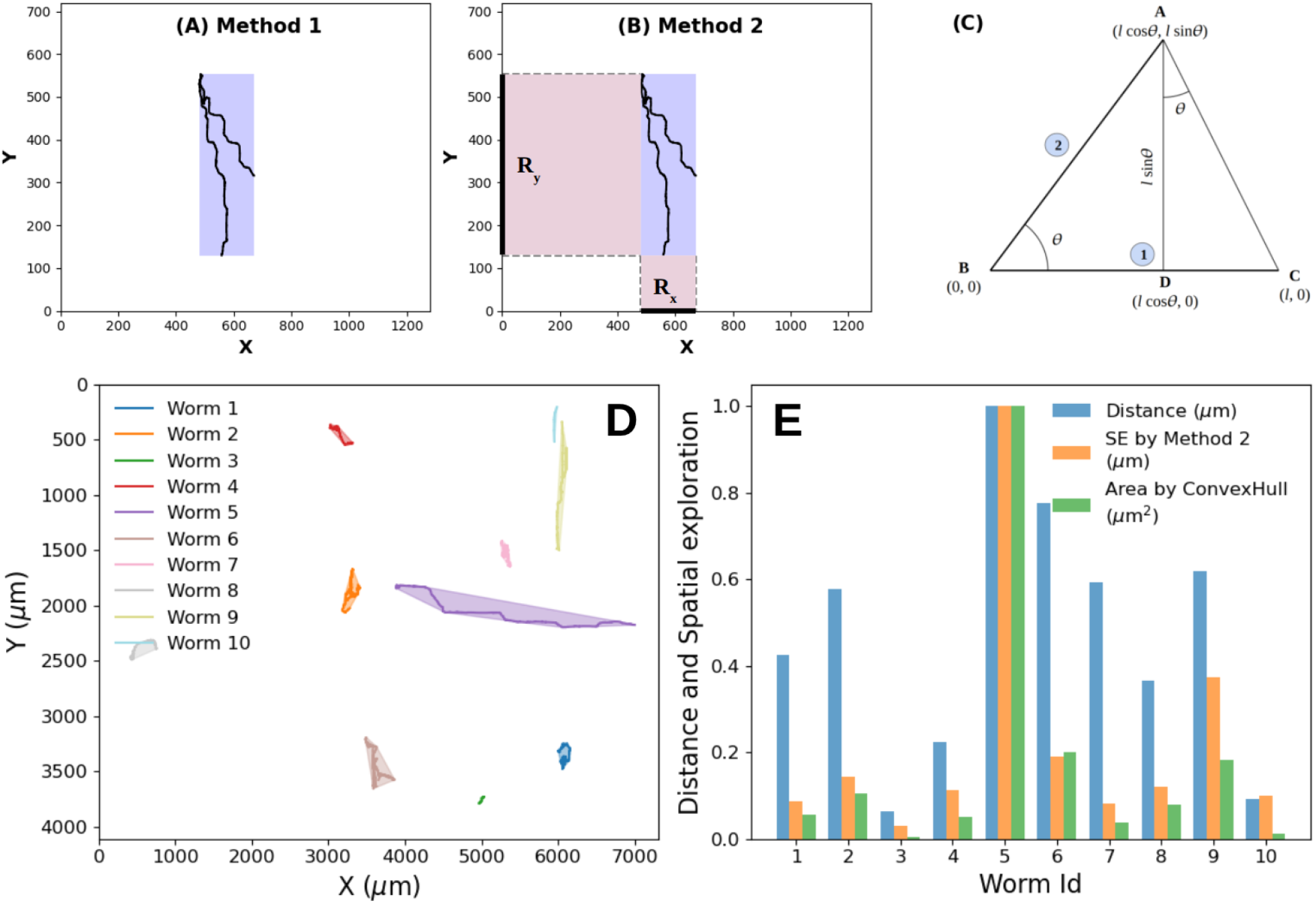
Measurement of regions explored by *C. elegans* during their locomotion. (A) Method 1 calculates the area of the rectangle covering the trajectory. (B) Method 2 calculates the spatial exploration by measuring projections *R*_*x*_ and *R*_*y*_ on each coordinate axis. (C) Illustration of Method 2 being independent of the orientation of the trajectory. (D) Trajectories of multiple worms representing distances covered by each worm. The shaded regions show the area covered by the trajectories calculated using ConvexHull. (E) Quantification of distance (*µ*m), SE using method 2 (*µ*m), and area using ConvexHull (*µ*m^2^) corresponding to each trajectory in Fig. D. The quantities are normalised with respect to their maximum values. Sample videos and outputs are provided in the [Google Drive](https://drive.google.com/drive/folders/1HM9xl3cHDyL2Bt_Z4EwOE5S-Ynpt1HAH?usp=drive_link).

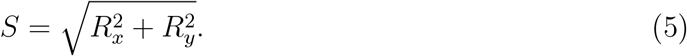

Here, *R*_*x*_ and *R*_*y*_ can be calculated from the trajectory of the worms using

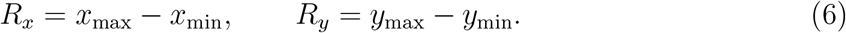

Now, let us see whether method 2 for measuring SE is independent of the orientation of the trajectory. To demonstrate that we can take two simple examples, as shown in Fig. 6C. Consider two trajectories 1 and 2, assumed to be the straight lines AB and BC (for simplicity), each of length *l*. Point B is assumed to be at the origin to further simplify the calculation. Trajectory 1 is parallel to the horizontal axis, whereas trajectory 2 is inclined at an angle *θ* with the horizontal axis. Now, *S* as per trajectory 1 is *S*_1_ = *l*, obtained from Eqs. (5) and (6). Area covered by trajectory 2 is 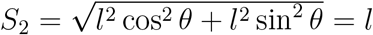. Therefore, both the trajectories have equal SE as per method 2, and are independent of their orientations. However, there are standard libraries for area measurement, which can potentially provide better quantification than method 2 does. One such library is the ConvexHull^7^ from SciPy, which draws a polygon around the trajectory by connecting the outermost points, and calculates the area covered by it. Therefore, the ConvexHull can be a good alternative to get a more accurate measure of the area covered by worms during their locomotion. Here we would like to emphasize (to avoid confusion) that, in this analysis, we refer to SE as the projections (*µ*m) measured by method 2, and the “area” is referred to as the area (*µ*m^2^) measured by ConvexHull. The reason for using dimensionally different quantities is that it is not known a priori which quantity can be a better representation to quantify the region explored by the worms.

Finally, to bring out the differences in the computed distance and spatial exploration, we have shown trajectories of multiple worms in Fig. 6D. The coloured shades around the trajectories represent the area covered by the worms according to the ConvexHull method. Each worm shows very distinct movement behaviours, and to characterize and quantify them, we have compared the distance, SE, and area of individual worms. The comparison is shown in Fig. 6E. Distances are calculated using the method described in Subsec. III B 3. There are wide variations among total distance travelled, SE measured by method 2 (*S*), and area measured by ConvexHull. Because of this, we normalize each of these quantities with respect to its maximum value, so that the results can be compared with each other. However, it should be noted that a valid comparison is not possible, as the quantities are dimensionally different. But our motive is to find the parameter which is a potentially better representation to quantify the extent of space a worm explores during its locomotion. From 6E, the Worm IDs 5 and 9 travel long distances, as shown by their trajectories. The corresponding SEs are also higher. But, for Worm IDs 1, 2, 6, 7 and 8, distances covered are high, but they explore relatively smaller regions, as their movements traverse shorter ranges. There are worms like Worm ID 4 and 10, which travel shorter distances as well as cover smaller regions. There are also worms like Worm ID 3, which do not show any movement. Now, based on these distances and SE measurements, we classify these locomotions qualitatively, which is shown in Table VI. Class I corresponds to the roaming state, whereas classes II and III can be identified as the dwelling movements. From Fig. 6D, it is evident that the ConvexHull method gives a more accurate measure of the area covered by the trajectories. But it can suffer from the same problem as method 1 does. For instance, the trajectory of Worm ID 9 in Fig. 6D is nearly vertical; therefore, the area measured by ConvexHull is very small. A careful observation will show that the area measured for worm ID 9 is approximately the same as that of worm ID 6, according to the ConvexHull method. But from the trajectories, worm 9 definitely explores more space than worm 6. Therefore, we conclude that ConvexHull gives a more precise quantification of the area covered by the trajectories, while method 2 is more robust for measuring spatial exploration, as it is independent of the shapes and orientations of trajectories.

**Table VI:** Classification of worms’ locomotion based on their distance and SE measurements as shown in Fig. 6E.

| Class | Distance | SE ($S$) | Worm ID | Locomotion type |
| --- | --- | --- | --- | --- |
| I | High | High | 5, 9 | roaming |
| II | High | Low | 1, 2, 6, 7, 8 | dwelling |
| III | Low | Low | 4, 10 | dwelling |
| IV | $\sim 0$ | $\sim 0$ | 3 | no movement |

#### 5. Orientation of motion

The nematodes *C. elegans* occasionally show reversals (switching direction of movement and moving in the reverse direction) when they come across any unfavourable situation in their surroundings. Worms can experience various kinds of environmental stimuli, such as chemical, mechanical, thermal or optical stress. The very first phenotypic response is to change their motion direction to get away from this stress. In many cases, this process happens systematically, by first reversing their motion, turning around, and then moving forward towards the favorable environment [79]. Understanding such behaviours is of immense importance for studying various signal-response dynamics and signalling pathways. Therefore, automated quantification of reversals is another step forward towards behavioural studies of *C. elegans*. Using the proposed method, we can determine the orientation along which a worm moves. When it reverses its motion, the orientation is changed by a large angular shift. By observing this shift, we can determine whether the worm has changed its orientation or not. The advantage of this method is that we need only the trajectory data to obtain the movement orientation, and it does not require any reference point. One can, if required, make a note of the initial orientation and infer the change in directionality from it. This means that if the worm is initially moving in the forward direction, a large angular shift would indicate that the worm has taken a reversal, and the exact moment of reversal can be accurately captured.

To evaluate the orientation, we first calculate the instantaneous speed of the worm at each frame. The instantaneous speed can be calculated as 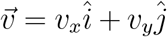, where *v*_*x*_ = *dx/dt, dx* = *x*_*i*_ − *x*_*i*−1_, and *dt* = *t/N, N* being the total number of frames. The magnitude of the velocity can be evaluated as 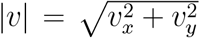, and the corresponding direction can be calculated from *θ* = tan^−1^(*v*_*y*_*/v*_*x*_). Evaluating this angle *θ* gives the orientation along which the worms move. If a worm undergoes any orientation changes, the velocity vector will change, and there will be a large shift in the orientation angle, as we describe in Fig. 7.

**Figure 7:**
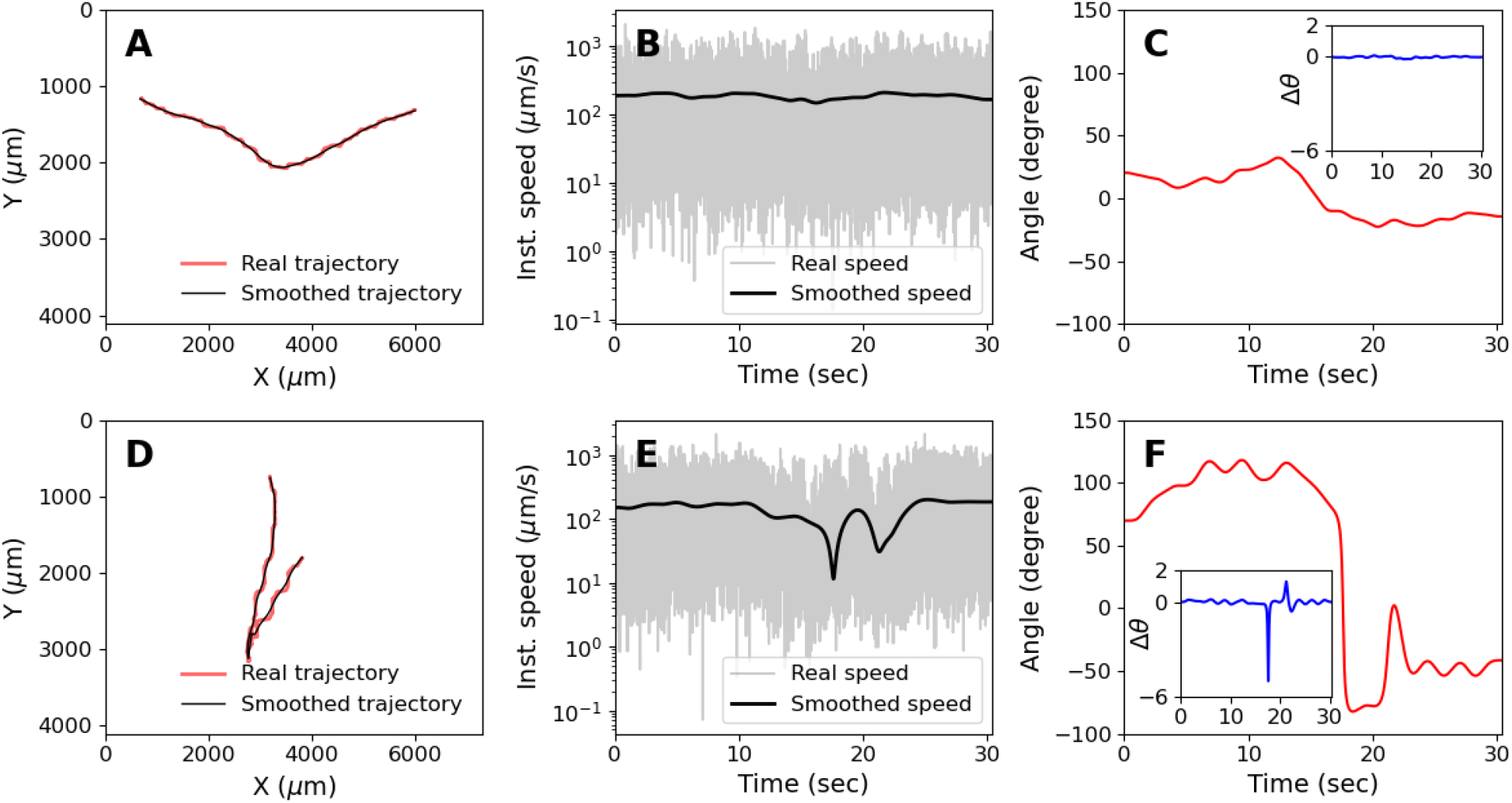
Orientation and instantaneous speed of *C. elegans* from their trajectories. (A) Trajectory showing no sharp change in orientation. The red curve shows the actual worm trajectory, and the black curve shows the filtered and smoothed trajectory. (B) Instantaneous speed (grey) and smoothed speed (black) after applying the Kalman filter. (C) Motion orientation, calculated from smoothed instantaneous speed. (D) Trajectory from a different video where the worm undergoes an orientation change characterized by a sharp edge. (E) Instantaneous and smoothed speed. (F) Change in orientation during locomotion. Orientation change happens twice, the first one at ~ 17 sec, and the second one at ~ 22 sec. Insets in Figs. C and F show the orientation change (Δ*θ* in degrees) at each frame. Video references are shared in the [Google Drive](https://drive.google.com/drive/folders/1i_PPpJpqyxJ-m72rK8t2Y0eAVLaToA08?usp=drive_link).

However, the instantaneous speed is extremely noisy, as shown in Figs. 7B and 7E in grey. Because of this, the orientation angle also fluctuates, which makes it difficult to capture the actual orientation change. Therefore, the first task while dealing with such noisy data is to smooth the speed profile. We can use the shifting-average technique, in which we consider a window of a certain width and calculate the average velocity within it. The window is then shifted to the next interval, and the same task is performed. This process is repeated till the last frame of the video. This method indeed smoothens out the fluctuations, but also introduces a time lag (proportional to the window size) between the actual and measured events in the video. Therefore, a more accurate technique is required which minimizes this lag. We used the Kalman filter^8^ to solve this issue, which efficiently reduces the fluctuations with negligible time difference.

To achieve smooth profiles of trajectories and speeds of the worms, we used a linear Kalman filter based on a constant-velocity state-space model independently to the *x* and *y*-coordinates of the trajectory. The hidden state vector is defined as **x**_*t*_ =[*x*_*t*_ *v*_*t*_]^⊤^, where *x*_*t*_ and *v*_*t*_ denote position and velocity of the worm at time *t* (similar notation follows for *y*-coordinates, throughout). The state evolution is then governed by

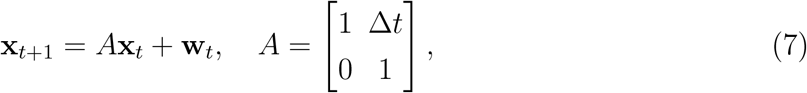

where Δ*t* is the time interval between consecutive frames, and **w**_*t*_ ~ *N*(0, *Q*) represents process noise. The process noise is a normally distributed random variable with zero mean and covariance *Q*, which has been chosen as 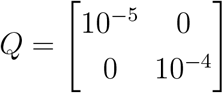, ensuring very small position uncertainty and slightly larger speed uncertainty, as the instantaneous speed is often more noisy. In the observation model, we provide trajectory as the raw input, and no input of speed is given. Therefore, positions are measured as

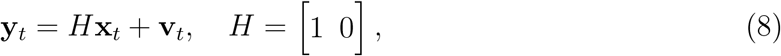

where **v**_*t*_ ~ *N*(0, *R*) is the observation noise, *i.e*., the noise in the input data. The observation noise is also a normally distributed random variable with zero mean and variance *R* = 0.2. Unlike the process noise, this value is relatively larger, reflecting higher uncertainty in detected positions. This ensures strong smoothing to follow the overall trend of the trajectory, without capturing minute details. The initial state was set as **x**_0_ = [*x*_0_ 0] ⊤, where *x*_0_ is the first observed position obtained from the trajectory of the individual worm (for faster convergence), and the initial speed is taken to be zero. State estimation was performed using the Rauch-Tung-Striebel (RTS) smoother (via forward-backward Kalman smoothing), which reduces temporal lag and provides smoothed profiles of both position and instantaneous speed as outputs over the entire trajectory [84]. Two independent Kalman filters were applied along individual components of the trajectory in order to estimate smoothed position and speed, *i.e*., 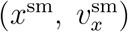 and 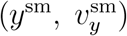. Finally, the orientation of motion is computed from the smoothed velocity components as 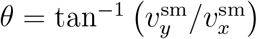, where 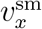 and 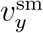 are the components of the instantaneous speed after smoothing. The orientation is unwrapped to avoid artificial jumps and to detect actual orientation changes.

Some relevant results are presented in Fig. 7, as outcomes from two different videos. The first row corresponds to a video in which the worm keeps moving in the forward direction, whereas, in the second row, the worm undergoes a direction change. Let us understand these two scenarios with the help of the proposed method. We first plot the trajectories in each case, which are shown in Figs. 7A and 7D. The actual trajectories (red) as well as the Kalman filtered trajectories (black), both are shown. The speed profiles are shown in Figs. 7B and 7E, in which the grey data represents the actual noisy instantaneous speed, and the thick black line is the smoothed speed after using the filter. The movement orientations are shown in Figs. 7C and 7F, which are computed from the smoothed trajectories using the proposed technique. In the first case, the worm does not change its movement direction, and no sudden jump in orientation angle is observed in Fig. 7C. Whereas, in the other case, the worm changes its locomotion direction from forward to reverse, and a large shift is observed at ~17 sec, as shown in Fig. 7F. The worm keeps moving in the reverse direction till ~22 sec, and changes its orientation again to start moving in the forward direction. The jumps in orientation (Δ*θ*) are shown as insets in Figs. 7C and 7F, in order to enable visualization of significant orientation changes. Interestingly, the smoothed speed in Fig. 7E drops to a lower value when orientation change happens. It makes sense because when the worm changes its movement direction from forward to reverse, it stops momentarily, which causes a reduction of the instantaneous speed. Similarly, when it changes the orientation again at ~22 sec, the speed drops again. These are very subtle effects of directionality change, but are captured efficiently by the algorithm.

#### 6. Forward-reverse locomotion

In the previous method, as explained in III B 5, we discussed the technique to detect any orientation change from the trajectory data. However, it cannot distinguish whether the motion is in the forward or reverse direction. Here, we describe a simple technique that can identify the forward and reverse movements with the help of keypoints. We constructed two vectors 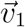 and 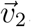, where 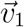 is the body vector representing the orientation from tail to head, and 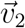 is the movement vector showing the locomotion direction. The orientation of both of these vectors can change as the worm moves, but 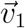 is always pointed from tail to head, while 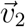 can change its orientation depending on the movement direction. Now, if the worm moves in the forward direction, both the vectors are pointed along the same direction (parallel). In contrast, during reversals, the vectors are pointed in the opposite direction (antiparallel). Some representative images are shown in Fig. 8A and 8B, where the directions of 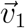 (green arrow) and 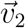 (blue arrow) are shown, corresponding to different locomotion states. To determine the parallel and antiparallel orientations between 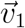 and 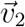, we first calculated the unit vectors 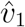 and 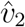, and took the dot product between them. In this way, we obtained 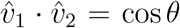, since the magnitudes of 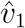 and 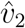 are unity. The values of the dot product can vary in the range [−1, 1]. Next, we applied the condition: if 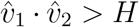, the motion is in the forward direction, and if 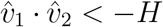, the motion is in the reverse direction, where, *H* is the threshold. Note that no locomotion phase is detected when 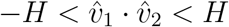. The quantities 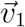 and 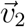 vary widely across frames; therefore, to reduce fluctuations, we used a temporal window, or BUFFER. The unit vectors are averaged over this window. At the beginning, the algorithm does not detect any direction, but once the buffer time is crossed, movement directions are estimated based on the orientations of 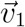 and 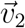. But there can exist instances where the bodies of the worms undergo compression and elongation while keeping the body stationary. This can lead to false detections of reversals. To remove these false positives, we employed the following rule in addition to the existing conditions. The body centroid of the worm must have moved by a certain distance in order for the movement to be considered valid. But this certain distance is highly subjective, and can vary from one experimental condition to another. Therefore, we needed a reference with respect to which the model could decide if a movement is valid or not. A natural reference is the length of the worm itself. So, the condition was applied as follows: if the movement (*d*) was larger than a certain fraction (*α*) of the total body length (*L*), *i.e*., *d > αL*, it would be considered a valid movement. Here, *d* is the distance travelled by the worm during a reversal, and *L* is the arc length of the worm’s body obtained by measuring and summing the distances between keypoints. If *d < αL*, the worm is considered stationary.

**Figure 8:**
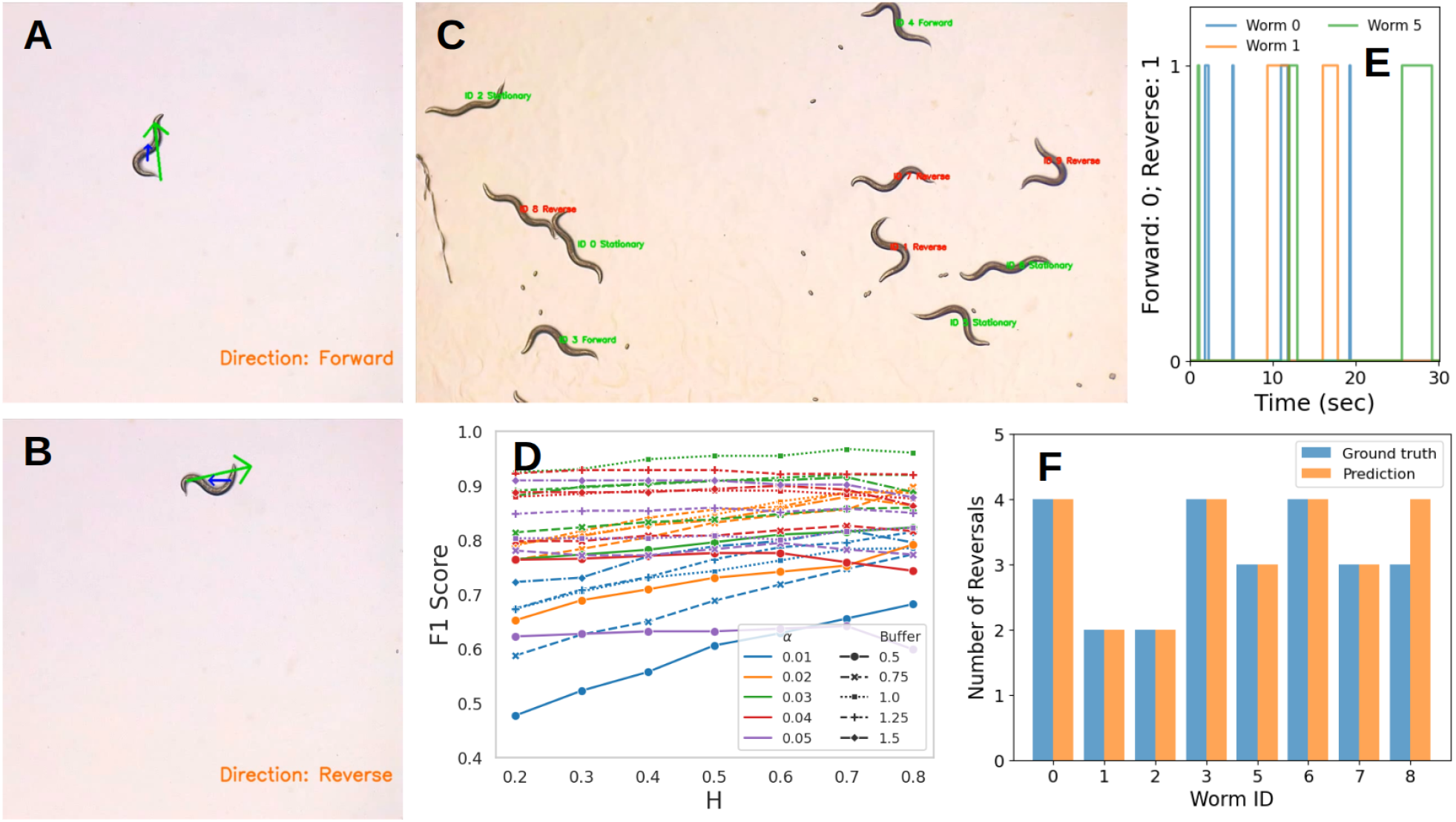
Detecting forward-reverse motion during worm movement. (A) and (B) show whether the worm is moving in the forward or reverse direction, depending on the body vector (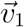, shown in green) and movement vector (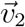, shown in blue). (C) Reversal detection on a video with multiple worms, in which the movement states are displayed. (D) Optimization of parameters for the detection of reversals. The detection accuracy is shown in terms of F1 score, corresponding to the angle threshold (*H*), body-fraction (*α*), and Buffer size. (E) Forward and reverse motions are shown by numbers 0 and 1, respectively, with time on the X-axis showing time instants of reversal events. It also shows the total number and the durations of each reversal event corresponding to each worm. (F) Number of predicted reversal events and comparison with ground truths. Video references are available in the Supplementary Materials (S4) and in the [Google Drive](https://drive.google.com/drive/folders/1xzgAVWeizEftUC316w83z8ZTroNr8sku?usp=drive_link).

Now, we have three parameters – the angle threshold (*H*) between 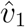 and 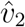 unit vectors, the buffer size, and the body length fraction *α*, which needed to be optimized before implementation in order to identify reversals with the highest possible accuracy. To do that, we prepared a dataset that contained manually annotated ground truth (GT) with the start and end times of each reversal event for every worm. We used a total of 5 videos of multiple worms, consisting of approximately 110 reversal events. For preparing the GTs, we have used only those worms that are visible in the entire video, so that we do not encounter any false positives during the optimization. The optimization was performed on 4 different videos, with the metrics TP, FP and FN evaluated with respect to the GT. TPs are those events for which the predicted intervals overlap with the GT intervals corresponding to the reversals of individual worms. FPs are those which are detected but are not present in GT, and FN are those which are in GT, but the model fails to detect them. Basically, FPs and FNs are those for which the intervals of prediction and GT do not overlap. With these parameters, we have calculated the precision and recall, and then calculated the F1 score, which represents the detection accuracy corresponding to each combination of these tunable parameters. A representative figure is shown in Fig. 8D, where the F1 score is plotted for different values of the parameters, represented by colours and symbols. From our analyses, the best performance was obtained for *H* = 0.7, BUFFER = 1 second (100 fps), and *α* = 0.03. Once the optimization was complete, we used these values and ran the algorithm on the test dataset. Testing was performed on a multiple-worm video, whose snapshot is shown in Fig. 8C, and the summary of total reversal counts for individual worms is shown in Fig. 8F. The frame-wise forward-reversals are shown in Fig. 8E, where reversals are indicated by 1. This jump from 0 to 1 indicates that the worm has changed it orientation from forward to reverse direction. It is also helpful to see which worm underwent reversal at which instance of time, and for how long it stayed in that state.

There are certain issues that need to be mentioned. The proposed algorithm is useful for detecting the directionality, but the exact instants at which transitions from one state to another happen is difficult to identify precisely, because of the presence of the temporal window used to reduce fluctuations. For more accurate time series detection, more precise filtering methods should be used.

#### 7. Quantifying omega turns

*C. elegans* often show different types of body bends, such as omega and delta turns, when they move. It is typically observed in stressed environmental conditions when the organisms tend to change their locomotion directions. That is the reason that these patterns are often associated with reversals [79, 85]. There is no universal method for detecting omega turns; however, several methods have been proposed in the past. For example, Gray et al. [79], in their experimental study, observed that during omega or delta bends, a worm reorients its body by more than 135^◦^. Following that, Huang et al. [85] proposed an algorithm based on keypoint detection, where they apply the condition that the bending angle between the lines connecting the head and mid-body, and the mid-body to the centroid, be less than 45^◦^. They have also conditioned that the length of these two connecting lines has to be in certain proportions to the body length for correctly identifying the start and end of the omega or delta turns. In a recent work, Suwazono et al. [86] proposed another algorithm that, during an omega bend, the head approaches its body, and based on that, they measured the distance between the head and any body segment to quantify such bendings.

Our proposed method is a combination of these two methods – as we have used the conditions on angle, as well as the body end-segment distances, to identify an omega or delta turn. The bending angle (*θ*) has been calculated between two lines connecting the head to the body centroid, and the body centroid to the tail. A representative image is shown in Fig. 9A, where the angle *θ* has been calculated between the two lines connecting the keypoints 0 to 5, and 5 to 10. The angle is calculated for all possible body shapes that appear in each frame, and we put the condition that if *θ < θ*_th_, then one can assume that the omega bend has formed. Here *θ*_th_ is the bending angle threshold, which we kept as a free and tunable parameter, to find the best value (or range of values) that gives accurate detection results. However, conditioning solely on the bending angle can lead to false positives, which requires us to incorporate additional conditions to get more accurate predictions. During the formation of an omega/delta turn, the head of the worm approaches the body, as described by Suwazono et al. [86]. When the bending is most prominent, the head segment becomes very close to its tail segment. Following this logic, we have calculated the proximity between the head and tail parts of the body in every frame. The proximity is calculated by measuring the distances between the first and last three keypoints, instead of measuring the head-tail distance only. A schematic is shown in Fig. 9A, in which the proximities between the keypoints 0, 1, 2, and 8, 9, 10, are measured, which we named as *d*_ht_. Therefore, *d*_ht_ is a 3 *×* 3 matrix with each of its elements representing the distance between the above-mentioned keypoints, and is calculated in every frame of the video. Now, during an omega or delta turn, the head and tail regions come very close to each other. To incorporate that, we apply the condition that if at least one element of the *d*_ht_ matrix becomes lower than a threshold, and the bending angle becomes less than *θ*_th_, then the model considers the shape to be an omega turn. Equivalently, if any one of these two conditions is violated, it can be considered as the end of the omega turn. Notice that for a valid omega turn, both conditions should match. This is necessary because sometimes worms tend to make a turn, but instead of completing the turn, they move in some other direction. Therefore, applying an “or” condition at the beginning would lead to false positives.

**Figure 9:**
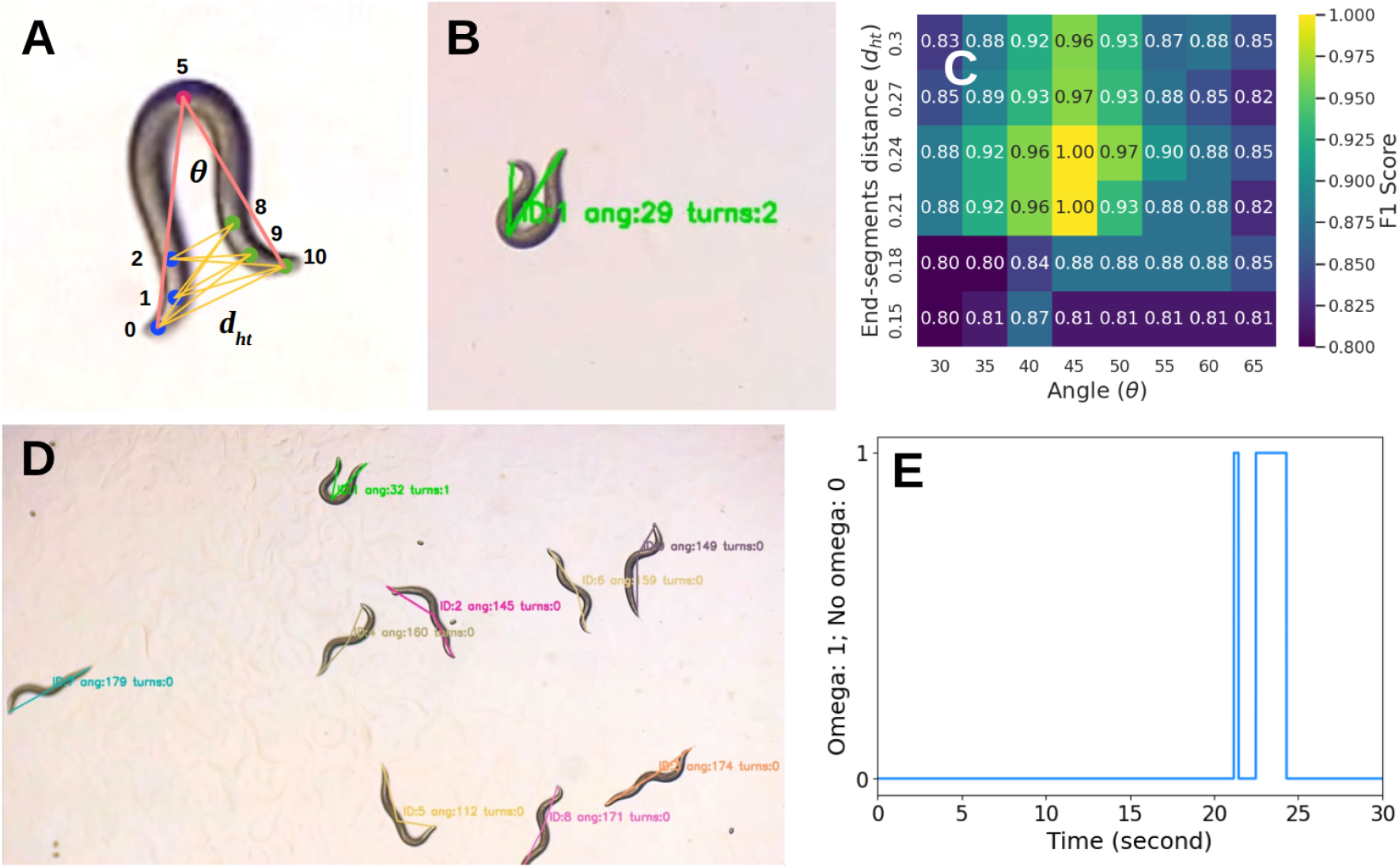
Detection of omega turns during locomotion of *C. elegans*. (A) Representation of the bending angle (*θ*) and the head-tail proximity (*d*_ht_) measurements. (B) Snapshot of a single worm making an omega bend, and detection of the individual turns from the video. (C) Optimization of threshold values for end-segment distances and the bending angle to achieve the best performance is evaluated using the F1 score. (D) Detecting and counting omega turns from videos of multiple worms. (E) Omega turns are denoted by 1; otherwise, it is zero. Therefore, a jump from 0 to 1 is observed when an omega turn is formed. Reference videos corresponding to D and E can be found in the [Google Drive](https://drive.google.com/drive/folders/1_X1iY7dTVIy27w5SpSmctsuAgNlNH3BC?usp=drive_link). The reference video for Fig. B is shared as a Supplementary Material (S5).

Once the conditions are set, the next step is to optimize the algorithm to find the threshold values for the distance and bending angles. We prepared a GT dataset containing omega turn events from different videos. Visually, it is difficult to identify the exact moments when an omega turn begins and ends. Therefore, we record the times at which an omega bend is visually most prominent. During detection, if an omega turn is predicted within a given window around the time mentioned in GT, the prediction is considered to be a TP, and if not, then it is a FN. Similarly, if an omega turn is predicted by the algorithm, but is not present in the GT, it is considered as FP. Using these values, we have calculated the precision, recall and F1 score for different values of bending angle and end-segments distance as thresholds. The optimization was performed on a subset of the entire dataset, and the corresponding F1 score is shown as a heatmap in Fig. 9C. The best prediction is obtained for the angle threshold of 45^◦^, which agrees with the previously reported value [79, 85], and a distance threshold ranging between 0.21 and 0.24. A snapshot of an omega turn in the test video is shown in Fig. 9B. A similar prediction on multiple worms video is shown in Fig. 9D. For visualization purposes, we have represented no-omega turns as 0 and the occurrence of an omega turn as 1; the corresponding result is shown in Fig. 9D. This makes it easy to visualize how many times and for what duration individual worms performed omega turns. It should be noted that the algorithm is suitable for identifying the omega turns. But the exact time for the start and end of an omega turn is subject to the conditions applied for detection.

#### 8. Multi-class object detection and counting

Simultaneous detection, tracking, and counting objects of multiple classes can be another useful assay for understanding animal behaviours during developmental stages. While working with a multi-class population, experimentalists may need to simultaneously detect, track, and count species. For example, counting the number of eggs can be valuable for understanding metabolic processes and reproduction. Similarly, we often perform experiments where worms of different ages (adult, L4, L3, *etc*.) co-exist. We may encounter situations where it is necessary to detect worms and eggs, or worms at different stages, simultaneously. This is beyond the capability of classical and other traditional DL models, as they are not optimized for such tasks.

Since this task involves only object detection, we prepare a model which is separate from the DPT. The training dataset was prepared, containing images of N2 adult worms and eggs on the same plate. In the dataset, some images were extracted from videos, and the rest were standalone images. To annotate images, we manually drew bounding boxes around the worms and eggs in the Roboflow framework. Two different classes were used – one for the worm, and another one for eggs. We had a total of 416 images, with 7554 annotations (5615 egg annotations and 1939 worm annotations). After annotations, we split the dataset following the same method that we used for the keypoints data as described in Sec. II D. This led to a split of 77:12:11 across the training, validation and test sets. Different augmentations have been applied, such as horizontal flip, rotation, shear, crop, grey scale and blur, which resulted in increasing the training images up to three times, while adding variability to the dataset. For training, we used different `YOLOv8` architectures for object detection, and training was performed on images of 768 × 768 resolution. The dataset has been provided in the [Google Drive](https://drive.google.com/drive/folders/1ayMr8-8SpxNPJe2DD3QNLO4JUcPIkUb5?usp=sharing). The rest of the training hyperparameters were the same as mentioned in Table II, except for the learning rate, which we fixed at 0.001. The batch size varied between 24 and 32, depending on the model architecture. The best performance was obtained from `YOLOv8x.pt`, which has precision = 0.967, recall = 0.949, mAP@0.5 = 0.982, and mAP@0.5:0.95 = 0.824, on the validation dataset.

The performance was verified on the test dataset, which contained 38 images with varying backgrounds and population densities. Sample examples are presented in Figs. 10A, 10B and 10C, which show detected worms and eggs on images with different backgrounds. Numbers of worms and eggs are counted manually as well as by model from each image in the test set, and the result is shown in Fig. 10D. In the figure, we have shown the predicted versus manually counted eggs, which shows very good correlation. The model shows good detection accuracy, even for larger populations, as can be observed from the figure. It is interesting to see that for large populations, where eggs are very close to each other, the model detects almost all the eggs. The incorrect detections happen mainly due to the image resolution or visibility of the eggs, but not due to the population size. But, for overcrowded conditions, the model can exhibit poor performance. The counting errors are shown in Fig. 10E, by calculating the differences between the predicted and manual counts with respect to the manual counting in each image. The error does not exceed by 2 units within the test dataset. The points on the horizontal dashed line correspond to the detections with no error. Similarly, we have plotted the error distribution to show weights corresponding to each error count, as shown in Fig. 10F. The vertical dashed line corresponds to the mean error over the whole test dataset. In addition to these, we have checked various counting metrics, such as the mean absolute error (MAE) and mean absolute percentage error (MAPE), which are calculated as

**Figure 10:**
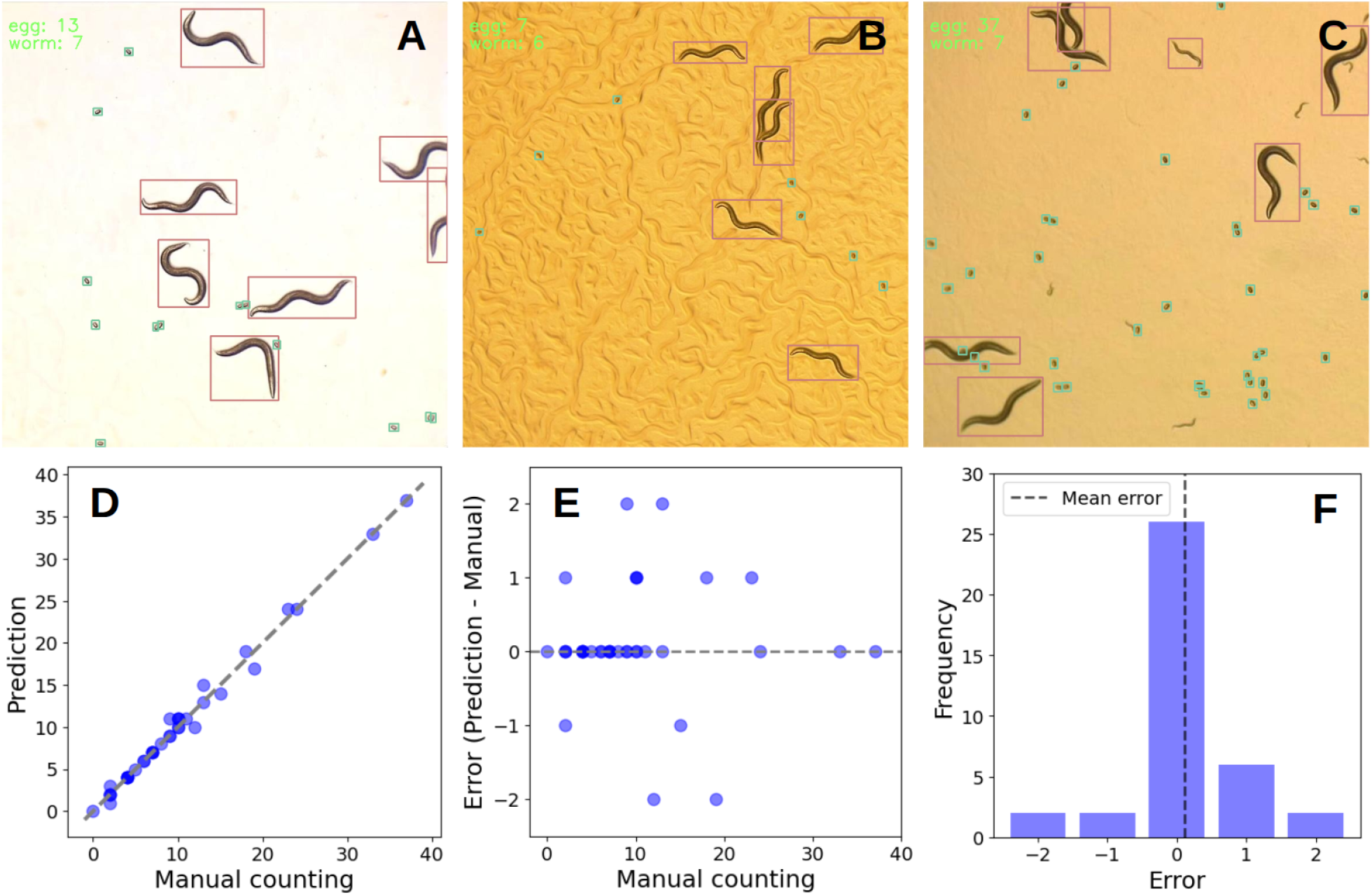
Detection and counting of multi-class objects. (A-C) Detection of worms and eggs in the test images. (D) Correlation plot to check model performance by comparing predicted versus manually counted eggs from the test images. The dashed line shows the *y* = *x* line. (E) Residual (error) of the predicted number of eggs from manually counted eggs from the images. (F) Counting error distribution to show how much weight each error holds. The dashed vertical line represents the mean prediction error over the complete test dataset. The outputs on the test images are shared in the [Google Drive](https://drive.google.com/drive/folders/1Z3F3V9JEszqEg8SoKRHPsA7Fu9Iuix0b?usp=drive_link).

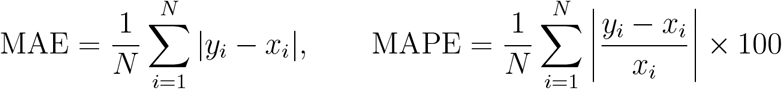

where *x*_*i*_ and *y*_*i*_ are the number of manually counted and predicted objects, respectively, from the *i*^th^ image, and *N* is the total number of images. MAE represents the average number of objects over- or undercounted compared to the manual counts, while MAPE is the average percentage relative error with respect to the manual counts. For our test set, the MAE is 0.42, and the MAPE is 5.56%, which indicates good accuracy while counting eggs and worms across varying populations. Despite very good accuracy, similar limitations as DPT hold for the detection and counting algorithm as well. For test images that are significantly different from the training data, the model can give poor performance, and model retraining might be required.

## IV. DISCUSSION

We propose Deep-Pose-Tracker for automated pose detection and tracking of body postures in *C. elegans*, an essential component in behavioural quantification. It performs bounding box and keypoint prediction on these nematodes, which is further integrated with several phenotypic analyses such as locomotion speed and spatial extent (area) measurements, motion orientation, forward-reverse motion detection, and identifying omega turns, *etc*. We have also enabled our model for eigenworms quantification, which is a useful technique to represent complex locomotion patterns in low dimensions. The model provides reliable tracking accuracy when dealing with moderately crowded populations, where worms can partially overlap with each other. The tracking metrics are shown in Table V, where the model maintains high tracking accuracies for a small population of worms, particularly when they partially or never collide with each other. As population size increases, overlaps and occlusions become more frequent, sometimes leading to identity switches and a corresponding reduction in tracking accuracy. In addition to tracking and pose estimation, we have designed a framework to detect objects of different classes and count them from videos and images.

Now, a natural question that can be asked is what are the advantages of using a YOLO based model, while there are several other classical image-processing and machine learning-based models that are particularly designed for *C. elegans*. The primary advantage lies in the underlying design of YOLO, as a unified deep learning framework for object detection and pose estimation. It is a popular deep learning model, that first appeared in 2016 [39], and gradually became one of the most popular DL models for object detection, tracking, classification, image segmentation, and pose detection, across diverse applications [33, 43, 47, 49–67]. Compared to other classical methods that rely on manual parameter tuning, YOLO models learn features directly from the dataset and yield predictions in a single forward pass. YOLO is known for fast inference and high accuracy, user accessibility, and, most importantly, the ability to detect objects of different classes simultaneously, within a single framework. Deep learning models usually require a large dataset to achieve good accuracy. But, in our case, we have used only 3018 images for training, and achieved *>* 99% detection and > 97% pose accuracy (mAP@0.5) on the validation dataset. This is possible because of the working principle of YOLO, which is based on the concept of transfer learning, in which the models are pretrained and fine-tuned on a large dataset. Transfer learning works on the same CNN architecture, consisting of a backbone for feature extraction and classification layers. The architecture is then trained on a large dataset (typically in the order of millions), and during the process, the feature extraction part of the model learns about the features and patterns present in an image. This trained model (known as the pre-trained model) can then be used to train on a relatively smaller dataset, while still providing strong performance compared to the models trained from scratch. During training these models, the pretrained weights are used and optimized, and that is the reason that the detection accuracy converges faster. The YOLO models for object detection are pretrained on the Microsoft COCO dataset^9^, containing 330k labelled images across 80 different classes. Similarly, the pose detection model was pretrained on the COCO-pose dataset^10^, which contains about 200k images of humans with 17 annotated keypoints across the body. The pretrained models for pose detection are then used to train on our own custom data containing *C. elegans* images, which resulted in strong detection and pose estimation performances.

We made a feature-wise comparison of DPT with previously existing models used for behavioural studies of *C. elegans*, particularly for pose detection. A summary is provided in Table VII. We describe this in detail in order to understand the capabilities and limitations of the DPT model, and the scenarios where it can be useful. Tracking worms in crowded environments or under noisy imaging conditions remains challenging for many automated tracking systems. In this work, we demonstrate that the DPT model can detect and track worms across different experimental conditions, including variable backgrounds and moderate crowding. The model can efficiently track partially overlapping worms; however, the performance can degrade under severe occlusions or heavy crowding. Several other locomotion features like eigenworms decomposition, detection of omega turns, detecting forward-reverse motion, locomotion orientation, *etc*., are integrated with the model, which are useful in behavioural studies. The model can detect worms irrespective of the presence of objects of different classes (*e.g*., eggs, or worms of different life stages) as demonstrated in Sec. III B 1. The YOLO object detection model is trained to detect multiple class objects simultaneously (like worms and eggs, or identifying worms at different stages), which is described in Sec. III B 8. This framework, in principle, can be used for other model organisms with appropriate training. Therefore, the model provides a unified framework for high-throughput analysis by integrating pose detection, tracking, and behavioural quantification within a single pipeline. The applicability of the model on a new dataset depends on the similarity of the imaging conditions, and may require retraining and fine-tuning based on the new dataset.

**Table VII:** Feature-wise comparison of DPT with pre-existing models for pose detection of *C. elegans*. “✓” represents the parameters that can be directly computed from the model as an output, whereas “×” shows the quantities that cannot be measured for a model. Further, there are some parameters which are not the direct outputs of a model, but can be measured as a post-analysis. Those are marked with “∗”. *Abbreviations* – DPT: Deep-Pose-Tracker, TT: Tierpsy Tracker, WP: WormPose, DT: DeepTangle, DTC: DeepTangleCrawl, DWT: Deep-Worm-Tracker.

| Parameters | DPT | TT<br>[24] | WP<br>[30] | DT<br>[32] | DTC<br>[36] | DWT<br>[33] |
| --- | --- | --- | --- | --- | --- | --- |
| Pose detection | ✓ | ✓ | ✓ | ✓ | ✓ | × |
| Multiple worms | ✓ | ✓ | × | ✓ | ✓ | ✓ |
| Head and tail identification | ✓ | ✓ | ✓ | × | × | × |
| Eigenworm quantification | ✓ | ✓ | * | * | * | × |
| Segmentation | × | ✓ | × | × | × | ✓ |
| Speed measurement | ✓ | ✓ | ✓ | * | ✓ | * |
| Spatial exploration | ✓ | * | * | * | * | * |
| Orientation of motion | ✓ | ✓ | ✓ | * | * | * |
| Detection of forward-reverse | ✓ | ✓ | * | × | × | × |
| Detecting omega turns | ✓ | ✓ | * | × | ✓ | × |
| Detect and track multiple class | ✓ | * | × | × | × | ✓ |
| Crowded environment | × | × | × | ✓ | ✓ | × |

Detection and tracking of worm body postures is an important aspect of studying animal behaviours. *C. elegans* shows a variety of body shapes and movement patterns under different environmental conditions, and understanding these is important for investigating neuronal circuits and signalling pathways. The DPT model enables automated pose detection of multiple *C. elegans* as shown in Fig. 3. Several efforts have been made earlier to estimate the animal poses [13, 25, 27, 30–33, 87–89]. However, a few of these can explicitly identify the heads and tails, particularly for *C. elegans* [24, 30]. One such model is Wormpose, which can estimate the body skeletons even for complex shapes involving overlap and self-occlusion, and can predict head and tail orientations [30]. It uses a probabilistic approach to estimate the head position and then incorporates a time series analysis across frames to validate the prediction. The detection is then evaluated over a separate dataset to verify if the detection is appropriate or not. The DPT model, on the other hand, uses a data-driven approach for head and tail identification without any post-processing. It learns feature representations directly from the image pixels during training, and uses those learned patterns to predict body keypoints in each frame during inference. Therefore, it works well on videos as well as on images. Wormpose particularly focuses on pose estimation of single worms, and are not explicitly designed for multi-worm tracking. In contrast, the DPT model is suitable for both pose detection and tracking of multiple worms within a single framework. It further addresses behavioural analyses such as locomotion speed and orientation, eigenworms decomposition, forward-reverse motion detection, omega turns *etc*. However, DPT does have some limitations as well. Since pose detection is performed independently in each frame, it does not carry any previous information for consistent head-tail identification, which can lead to occasional misidentification. Further, even though the model shows good accuracy on the validation dataset, the performance may degrade when the model is used for pose detection on images that are substantially different from the training data in terms of background, intensity or the resolution of the image, and where the heads and tails are not easily distinguishable. In such cases, the model needs to be retrained on the representative dataset.

A common challenge in deep learning models is the requirement of sufficiently large annotated datasets for training. To obtain the labelled data, one has to carry out manual annotations, which is typically time-consuming and labour-intensive. This can be circumvented by using synthetic data, which has previously been adopted and provided good performance on real videos or images [30, 35]. In addition, training deep learning models typically requires GPU-accelerated hardware, which is not always feasible. Cloud-based computations can be an alternative in such scenarios. However, once training is performed on a representative dataset, these models provide reliable predictions in image and video analysis tasks. The DPT model performs efficiently with multiple objects; however, tracking accuracy may decrease in crowded environments, where worms overlap heavily with each other very frequently. In our study, the model maintains good performance for moderate populations. Similar challenges may also arise in pose detection, as it requires good resolution for reliable predictions of heads and tails. If the features of heads and tails are not properly visible, or the worm makes highly coiled or complex body shapes, prediction errors can happen. The detection accuracy may potentially be improved by increasing the dataset size with more diversity in images, particularly by including data with more coiled and complex body shapes and challenging imaging conditions.

Future work can be done focusing on improving model performance in challenging situations, such as highly complex or coiled body shapes. One can incorporate temporal consistency across frames, which can help in maintaining the head-tail identity, that improve model performance and robustness.

## V. CONCLUSIONS

We present Deep-Pose-Tracker (DPT) for tracking and pose detection of *C. elegans*, essential for studying behaviours under varying experimental conditions. The YOLO-based model is trained on a custom dataset containing images of *C. elegans* with annotated keypoints to perform pose detection on videos and images. In addition, DPT is integrated with several post-detection analyses, such as estimating locomotion speed, eigenworm quantification, movement orientation, area explored, detecting reversals, and omega turns. DPT provides pose detection and tracking outputs, and is able to detect complex poses of multiple worms in challenging imaging conditions. The model is also reusable for different imaging setups after being trained on the representative dataset. We have also designed a YOLO-based counting framework for simultaneous detection of objects from different classes, which is essential in the study of animal behaviours and development. Therefore, the proposed framework includes a wide range of features that are useful in behavioural analysis and are important in the study of neuroscience, animal behaviours and signalling pathways.

## SUPPLEMENTARY MATERIALS

**S1 Movie:** Multi-worm pose detection with predicted keypoints and head-tail labels overlaid on the original frames.

**S2 Movie:** Another example of multi-worm pose detection with different background.

**S3 Movie:** Tracking multiple worms using YOLO and ByteTrack/BoT-SORT.

**S4 Movie:** Detection of forward-reverse orientation during locomotion.

**S5 Movie:** Detection of omega turns during locomotion.

## DATA AND CODE AVAILABILITY

All the codes, trained weights and dataset used in this study are publicly available on GitHub at: https://github.com/cebpLab/Deep-Pose-Tracker, with instructions for installation, training and deployment. To support long-term reproducibility, we provide a versioned release containing the details of the work, archived on Zenodo with DOI: https://doi.org/10.5281/zenodo.20035992.

## VI. ACKNOWLEDGEMENTS

We acknowledge Ultralytics for YOLO, a state-of-the-art Computer Vision model with various essential features, which is completely open-source. We thank Roboflow and CVAT for providing the annotation and data augmentation platform.

## VII. AUTHOR CONTRIBUTIONS

D.S. conceived the project, designed the methodology, developed the model (including training and validation), performed data analysis, and wrote the manuscript. S.C. and D.V. performed biological experiments, carried out the data acquisition, and provided biological interpretation of the results. A.G.R. helped with data acquisition and biological insights into the results. R.S. conceived and supervised the project, contributed to conceptualization and interpretation, and critically revised the manuscript for intellectual content.

## Footnotes

1 https://docs.ultralytics.com/

2 https://roboflow.com/

3 https://docs.ultralytics.com/tasks/pose/#models

4 https://www.cvat.ai/

5 https://pypi.org/project/motmetrics/#References

6 https://docs.ultralytics.com/modes/track/

7 https://docs.scipy.org/doc/scipy/reference/generated/scipy.spatial.ConvexHull.html

8 https://pypi.org/project/pykalman/

0 https://cocodataset.org/#home

10 https://cocodataset.org/#keypoints-2017

